## Supplementary material for "Plant community stability is associated with a decoupling of prokaryote and fungal soil networks"

**Supplementary methods**

*Seed collection and sowing*

Seeds of most of the plant species were collected from the natural grassland field site and a further 5% were obtained from nearby seed production fields (Planta Naturalis, Markvartice, Czech Republic). Seeds of all plant species were sown simultaneously. For this, seeds were placed on the soil surface and gently pressed into the soil to avoid seeds from blowing away[1].

*Soil chemical analyses*

All soil was sieved on a 2 mm mesh an thoroughly mixed. Plant available nitrogen (N) (mg $kg^{-1}$ dry soil) was determined by adding 50 mL of 0.5 M $K_2SO_4$ to 5 g of fresh soil, shaking for 30 min and filtering the soil out. $NO_3^-$, $NH_4^+$ and $NO_2^-$ concentrations in the filtrate were determined by Flow Injection Analysis (QuickChem 8000 FIA; Lachat Instruments, Loveland, CO, USA). Plant available P was determined following Olsen et al[2]. In brief, 5 g air dried soil was extracted with 50 mL of 1 M $NaHCO_3$ adjusted to pH 8.5 with addition of activated carbon to eliminate discoloration resulting from humic acid release. The solution was shaken for 2 h and soil was filtered out. Available P in the filtrate was determined by the Olsen photometric method (ATI Unicam UV 400/VIS Spectrophotometer at 630 nm)[2]. K was determined in 5 g air dried soil by shaking with 50 mL Mehlich II soil extraction solution (Hach Lange GmbH, Düsseldorf, Germany) for 30 min. Soil was filtered out and and Mg, Ca, and K were measured in the filtrate using atomic absorption spectrometry (ContrAA 700 with $C_2H_2$-air flame for Mg and K, and $C_2H_2$-$N_2O$ for Ca; Analytik Jena GmbH, Jena, Germany). Exchangeable pH was measured in a solution of 5 mL in 25 mL 0.1M KCl shaken for 30 min (WTW Multilab 540; Xylem Analytics, Weilheim, Germany). Total N, C and organic C were determined in dried soil ground to <0.1 mm particle size using combustion analyses (FLASH 2000 CHNS/O Analyzer; Thermo Fisher Scientific, Waltham, MA, USA).

*Soil bacterial and fungal biomass*

Soil bacterial and fungal biomass was determined using PLFA and NLFA analysis following García-Sánchez et al[3]. In short, 1 g of fresh soil taken from the mixed soil cores was freeze-dried in a chloroform-methanol-phosphate buffer (1:2:0.8, v/v/v)[4]. Lipids were fractioned into polar lipids (PLFAs), glycolipids and neutral lipids (NLFAs), using an extraction cartridge (LiChrolut Si-60; Merck KGaA, Darmstadt, Germany) and subjected to alkaline methanolysis[5]. Following Sampedro et al[6], free methyl esters of PFLAs and NLFAs were analysed by gas chromatography-mass spectrometry (450-GC with 240-MS IT Mass Spectrometer; Varian Medical Systems Inc., Palo Alto, CA, USA). Total microbial biomass was estimated by the sum of all PLFA contents. Bacterial biomass was based on the summed

PLFA contents i14:0, i15:0, a15:0, 16:1w5, 16:1w7; 16:1w9, 10Me-16:0, i16:0, i17:0, a17:0, cy17:0, 17:0, 10Me-17:0, 18:1w7, 10Me-18:0 and cy19:0, and actinobacterial biomass based on the summed contents 10Me-16:0, 10Me-17:0 and 10-Me18:0. Gram-positive and gram-negative bacterial were quantified based on i14:0, i15:0, a15:0, i16:0, i17:0, a17:0 and 16:1w7, 16:1w9, 18:1w7, cy17:0, cy19:0, respectively. Fungal biomass was quantified based on PLFA content 18:2w6,9[5] and NLFA 16:1w5 was used as a marker for AM fungi[7].

*16S and ITS amplicon sequencing*

All frozen soil samples (250 mg each, in duplicates for each sample) were homogenized and lysed in PowerBead Pro Tubes (Qiagen, Germany) on a Vortex adapter. Subsequently, DNA was extracted using the DNeasy PowerSoil Kit (Qiagen, Germany) according to the manufacturer's instructions and eluted in 50 µl of elution buffer. The fungal internal transcribed spacer of the rDNA (ITS2 rDNA) was amplified using primers gITS7ngs[8] and ITS4[9]. The bacterial 16S rRNA gene (V4 region) was amplified from the same DNA extracts using primers 515f and 806r[10]. All primers were tagged with sample-specific barcodes of 10-12 bases. PCR mix was performed in the total volume of 15 µl and contained 0.07 U Thermo Scientific™ *Taq* DNA Polymerase, 10x PCR Buffer, 2.5 mM $MgCl_2$, 20 µg BSA (all Thermo Fisher Scientific, Waltham, Massachusetts, USA), 0.3 mM each dNTP, 0.3 µM of each primer and 1 µl of the DNA extract. Thermocycling conditions were 94 °C for 4 min, 25 cycles of 94 °C for 45 s, 52 °C for 60 s and 72 °C for 35 s, followed by 10 min at 72 °C. Each DNA extract was amplified in duplicate. PCR products were visualized on a 1% agarose gel. The pooled duplicates were purified through columns with the QIAquick PCR Purification Kit (Qiagen, Hilden, Germany) according to the manufacturer's protocol and eluted into 20 µl of $ddH_2O$. DNA concentrations of the amplicon pools were quantified using a Qubit 2.0 Fluorometer (Thermo Fisher Scientific) with High Sensitivity Assay Kit. The purified amplicons were pooled in equimolar ratios. Both negative PCR controls (with $ddH_2O$ instead of a template) were processed in the same way as the experimental samples and included into the sequencing library, together with sixty fungal and sixty bacterial amplicons. The library was sequenced on an Illumina MiSeq instrument (2 × 250 bp) (SEQme, Dobříš, Czech Republic).

*Structural equation models*

To keep the number of potential pathways relative low and avoid spurious effects occurring due to correlating exogeneous variables, we first calculated three base models for each soil origin[11]. These three base models captured effects of the plant community onto soil chemical changes after the 13th growing season of (a) the plant community in the year of sampling (aboveground productivity, plant diversity and plant compositional DCA axes 1-3), (b) overall effects of the plant community from the past (initial invasion effect on plant diversity, aboveground productivity and plant diversity trajectories in time), and (c) plant compositional effects from the past (invasion effect size on plant compositional DCA axes 1-3, plant compositional DCA axes 1-3 trajectories in time). All base models included the same soil chemical parameters: total soil N, organic C and pH, and plant available P, $NO_3^-$, $NH_4^+$, $NO_2^-$ and belowground productivity. The three base models were then combined. Each microbial network cluster (summed relative reads per 16S or ITS cluster from the calculated co-occurrence networks per soil origin), bacterial, fungal and microbial biomass (PLFA/NLFA analyses) as well as alpha-diversity indices and average SI of the microbial communities were ran through the SEM model as the final parameter to be estimated. Per run, one microbial parameter was considered, which could be affected either directly by the plant community parameters or indirectly via soil chemical changes. Each run, a backward stepwise elimination procedure to consecutively remove non-significant pathways was followed in the same way as performed for the base models[12]. All microbial variables not following a normal distribution were ln- or sqrt-transformed.

*References*


1. Münzbergová, Z. Seed density significantly affects species richness and composition in experimental plant communities. *PLoS One* **7**, (2012).

2. Olsen, S., Cole, C., Watanabe, F. & Dean, L. Estimation of available phosphorus in soils by extraction with sodium bicarbonate. in *USDA Circular Nr 939* (US Gov. Print. Office, 1954).

### Supplementary figures

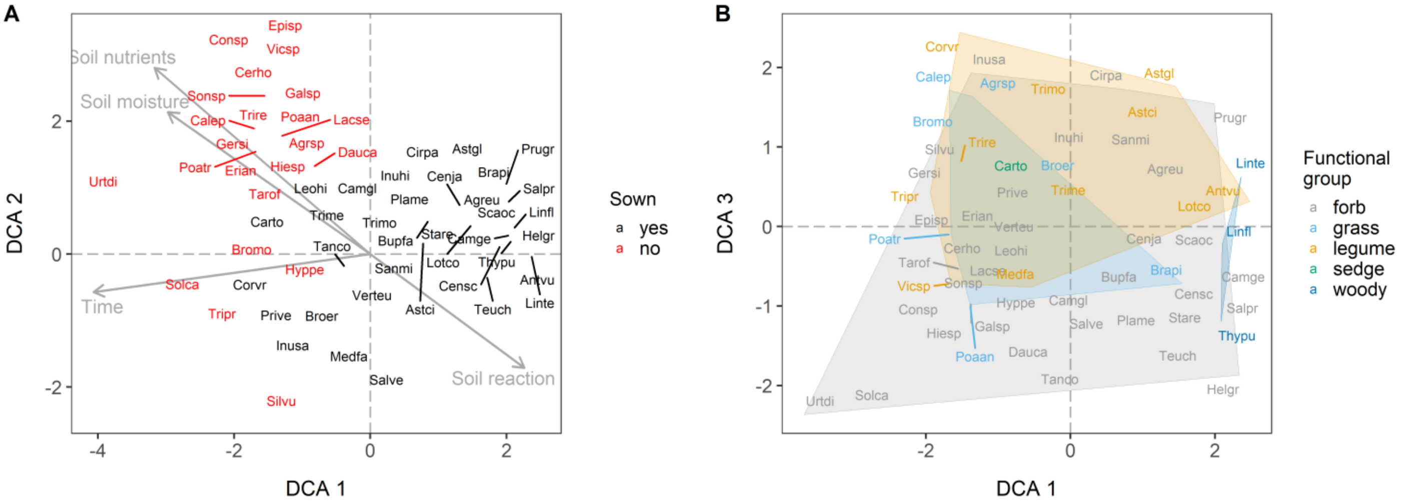


**Figure S1** Plant species position on detrended correspondence analysis (DCA) axes (A) 1 and 2, and (B) 1 and 3 (n = 59 plant species; both soil origins included). In A, arrows show a passive overlay of significant ($p < 0.05$) plant factors with 'Time' indicating the average time period (year between 2007 and 2019) that plant species were present in the communities, and 'Soil nutrients', 'Soil moisture' and 'Soil reaction' referring to plant species ecological optima based on Czech Ellenberg indicators. Plant species abbreviations shown in black are sown species, while species in red invaded the communities. In B, hulls indicate the section of the DCA axes where plant species of five functional groups occur and shows that DCA 3 separates communities based on legume cover. For plant species abbreviations, see Table S9.

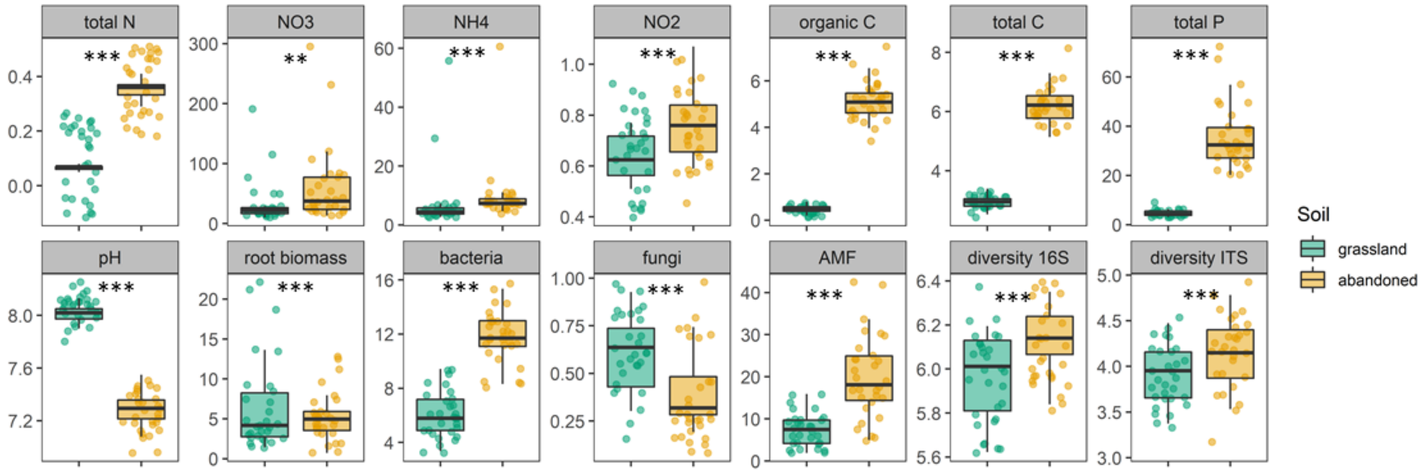


**Figure S2** Soil chemical and microbial properties after the 13th growing season in communities grown on natural grassland (green) and abandoned arable (yellow). Total N, total C and organic C are given in percentage; $NO_3^-$, $NH_4^+$, $NO_2^-$, K and P, as well as bacterial, fungal and AMF biomass are given in mg · $kg^{-1}$ dry soil; root biomass is given in gram, and for 16S and ITS diversity the Shannon diversity index is shown. Averages ± SE (*n* = 30), asterisks indicate significant differences between natural grassland and abandoned arable soil. Significance codes: *** $p < 0.001$; ** $p < 0.01$.

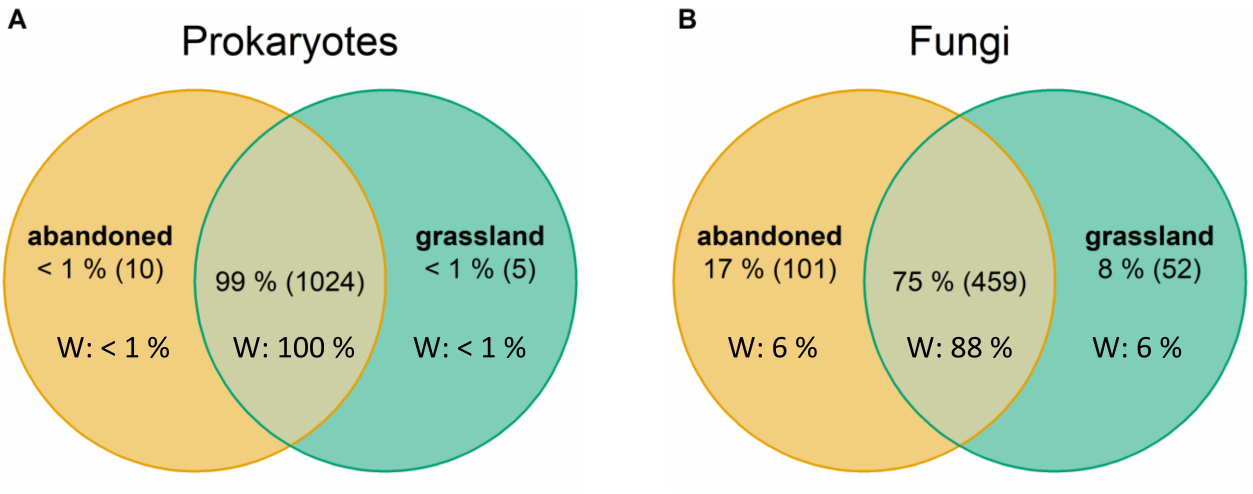


**Figure S3** Venn diagram showing the overlap in (A) prokaryote and (B) fungal OTUs between natural grassland (green) and abandoned arable soil (yellow). In the top row, the number of OTUs in each compartment is indicated in percentages and, in brackets, as the total number. The bottom row shows the percentage of OTUs in each compartment weighted by the OTUs relative abundance (n = 30 per soil origin).

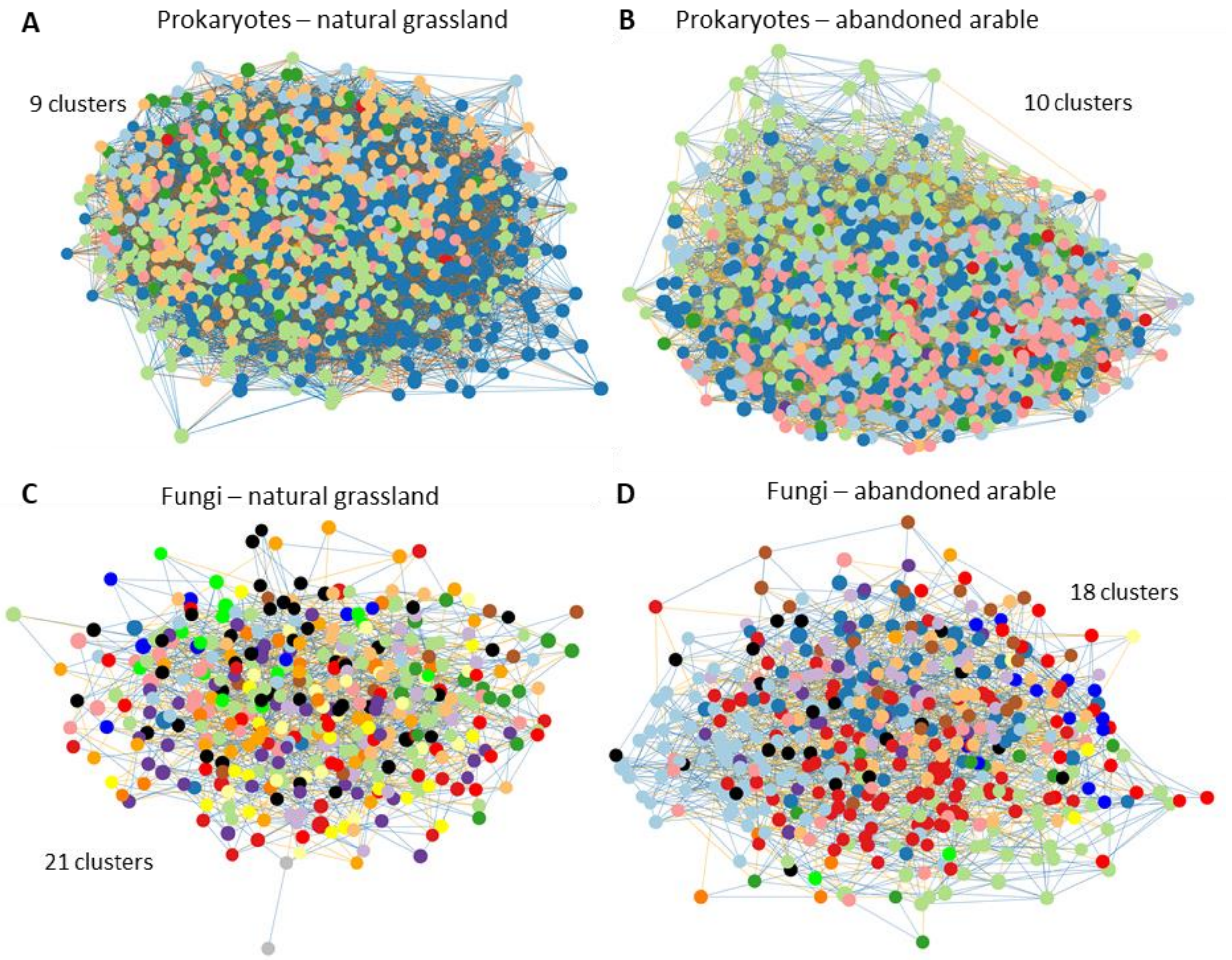


**Figure S4** Soil microbial co-occurrence networks of prokaryotes in (A) natural grassland soil and (B) in abandoned arable soil, and of fungi in (C) natural grassland soil and (D) abandoned arable soil. Each dot represents one OTU. Different colours indicate that OTUs belonged to different co-occurrence clusters within the network (*n* =30 per soil origin).

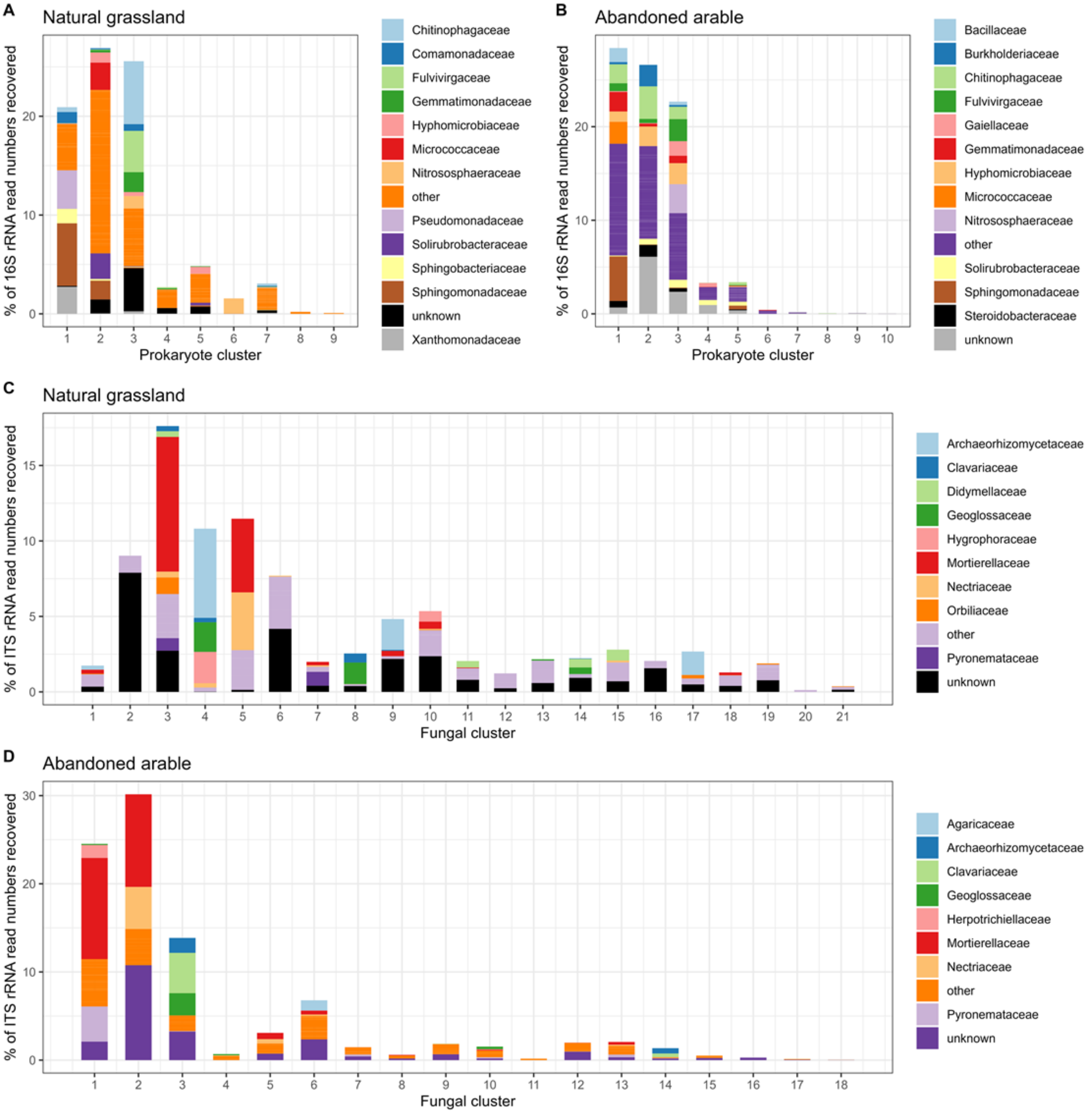


**Figure S5** Average percentage of 16S and ITS rRNA read numbers recovered per family level in prokaryote clusters in (A) natural grassland and (B) abandoned arable soil, and fungal clusters in (C) natural grassland and (D) abandoned arable soil. Microbial clusters were obtained from co-occurrence networks (Fig 3; S4). For 16S, families < 2% relative abundances are grouped in 'other', for ITS, families <1.5% relative abundances are grouped in 'other'.

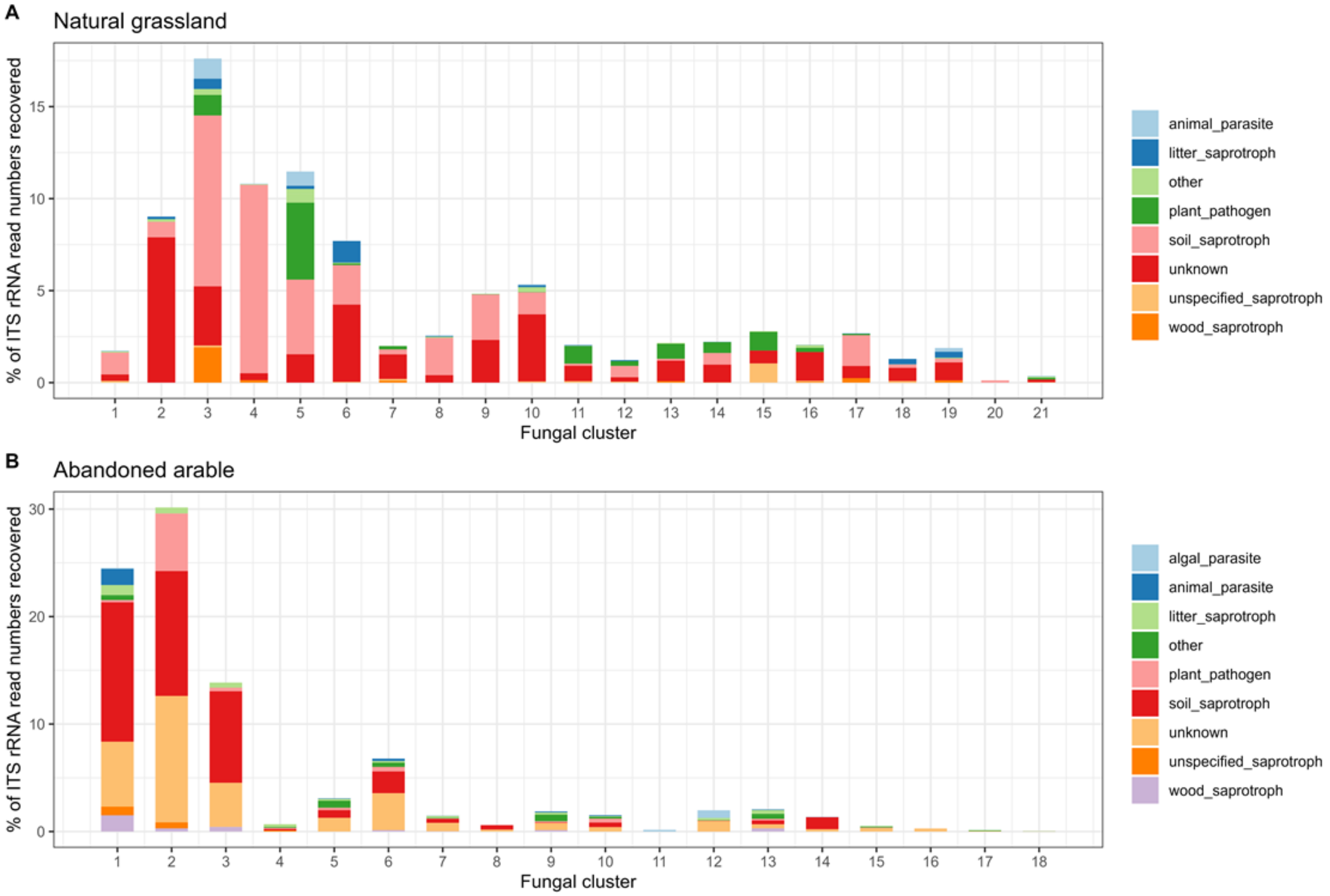


**Figure S6** Average percentage of ITS rRNA read numbers recovered per putative fungal trait over similarly responding fungal clusters obtained from co-occurrence networks in (A) natural grassland and (B) abandoned arable soil. Putative fungal traits < 1% relative abundances are grouped in 'other'. Fungal traits were obtained from the FungalTraits database (Põlme, S. et al. 2020. Fungal Divers 105).

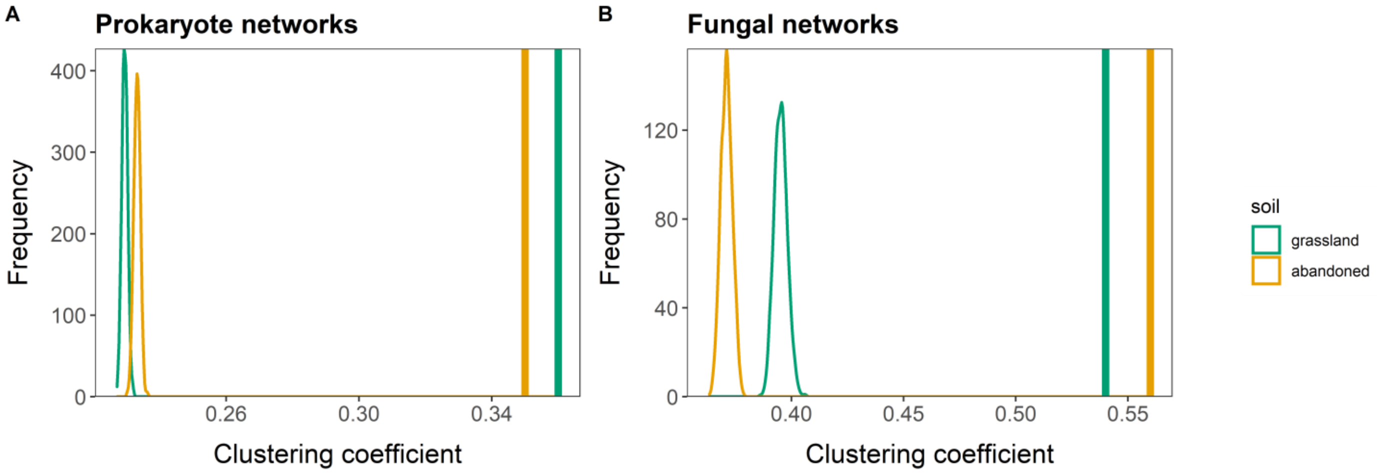


**Figure S7** Frequency distribution of clustering coefficients (modularity) in random networks of (A) prokaryote and (B) fungal co-occurrence networks in natural grassland (green) and abandoned arable soil (yellow). Vertical lines indicate the clustering coefficients of the real microbial networks. Random networks were created by rewiring the edges of the original networks while preserving the original networks degree distribution (1000 iterations). The clustering coefficients of the original networks never occurred in the rewired networks.

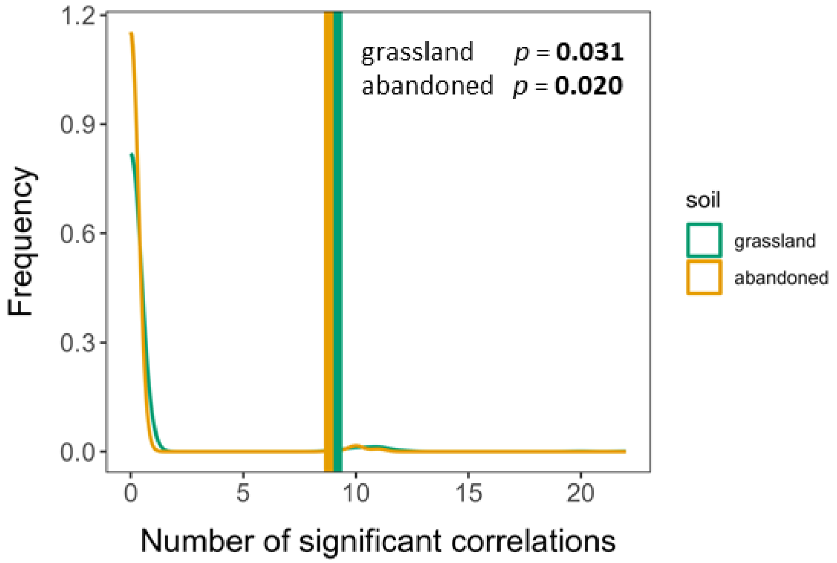


**Figure S8** Frequency distribution of the number of significant correlations between each prokaryote and fungal cluster in random co-occurrence networks in natural grassland (green) and abandoned arable soil (yellow). Vertical lines indicate the number of significant correlations in the real microbial networks (9 for both soils). Random networks were created by rewiring the edges of the original networks while preserving the original network degree distribution (1000 iterations).

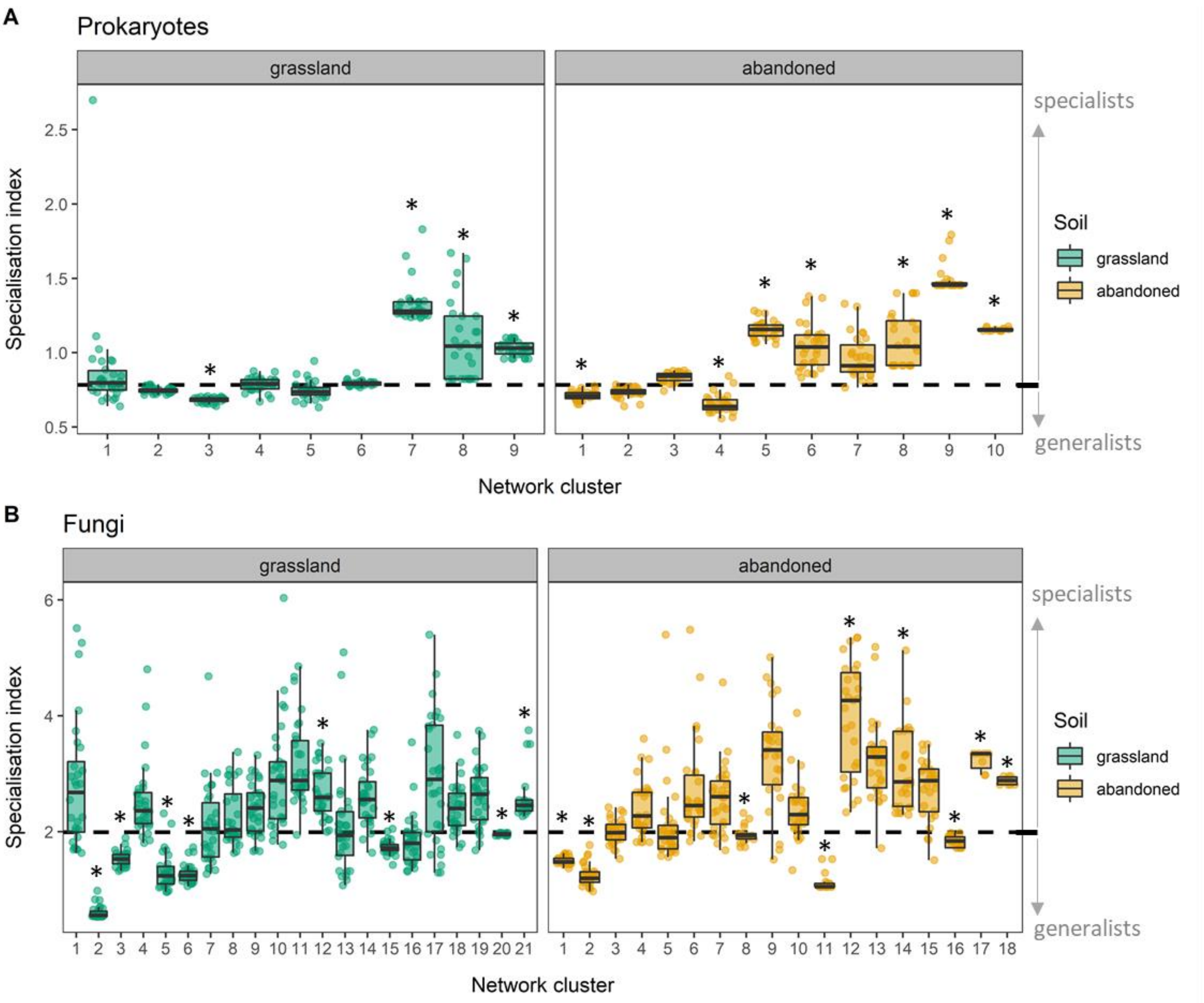


**Figure S9** Relative habitat specialisation (specialisation index; SI) of (A) prokaryote networks clusters and (B) fungal network clusters in natural grassland (green) and abandoned arable soil (yellow). The black, dashed line indicates the community-wide mean SI of the prokaryote and fungal community, respectively. Relative habitat specialist network clusters occur above the community-wide mean SI without overlap of the minimum distribution (25th percentile - 1.5 * interquartile rang; lower whisker). Relative habitat generalists network clusters occur below the community-wide mean SI without overlap of the maximum distribution (75th percentile + 1.5 * interquartile range; upper whisker). Relative habitat specialist and generalist clusters are indicated by an asterisk (n = 30 samples per cluster).

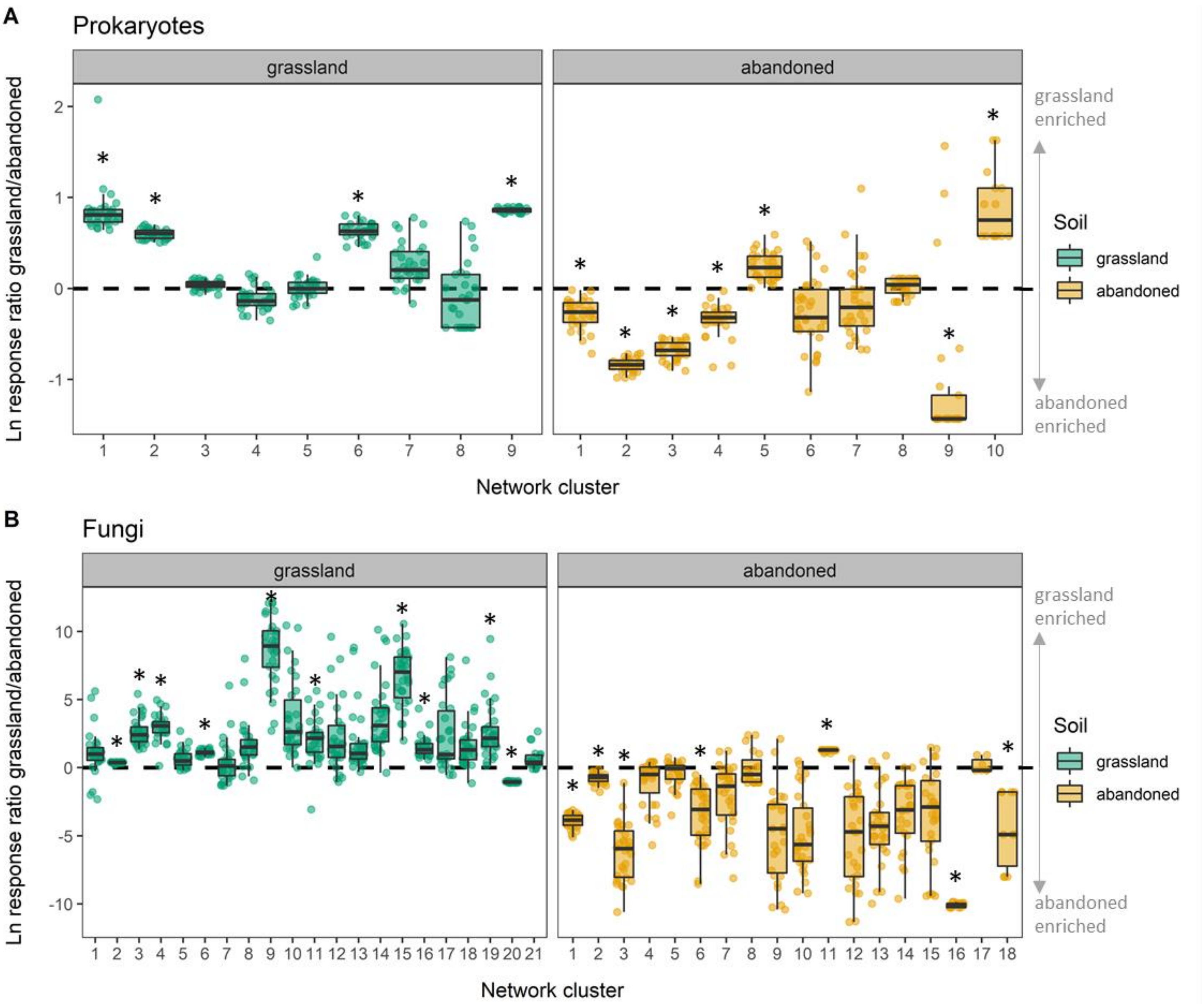


**Figure S10** Ln-response ratio OTUs abundances in natural grassland soil divided by abundances in abandoned arable soil for (A) prokaryote networks clusters and (B) fungal network clusters in natural grassland (green) and abandoned arable soil (yellow). Clusters enriched in natural grassland soil occur above 0 (dashed line) without overlap of the minimum distribution (25th percentile - 1.5 * interquartile rang; lower whisker). Clusters enriched in abandoned arable soil occur below 0 without overlap of the maximum distribution (75th percentile + 1.5 * interquartile range; upper whisker). Enriched clusters are indicated by an asterisk (n = 30 samples per cluster).

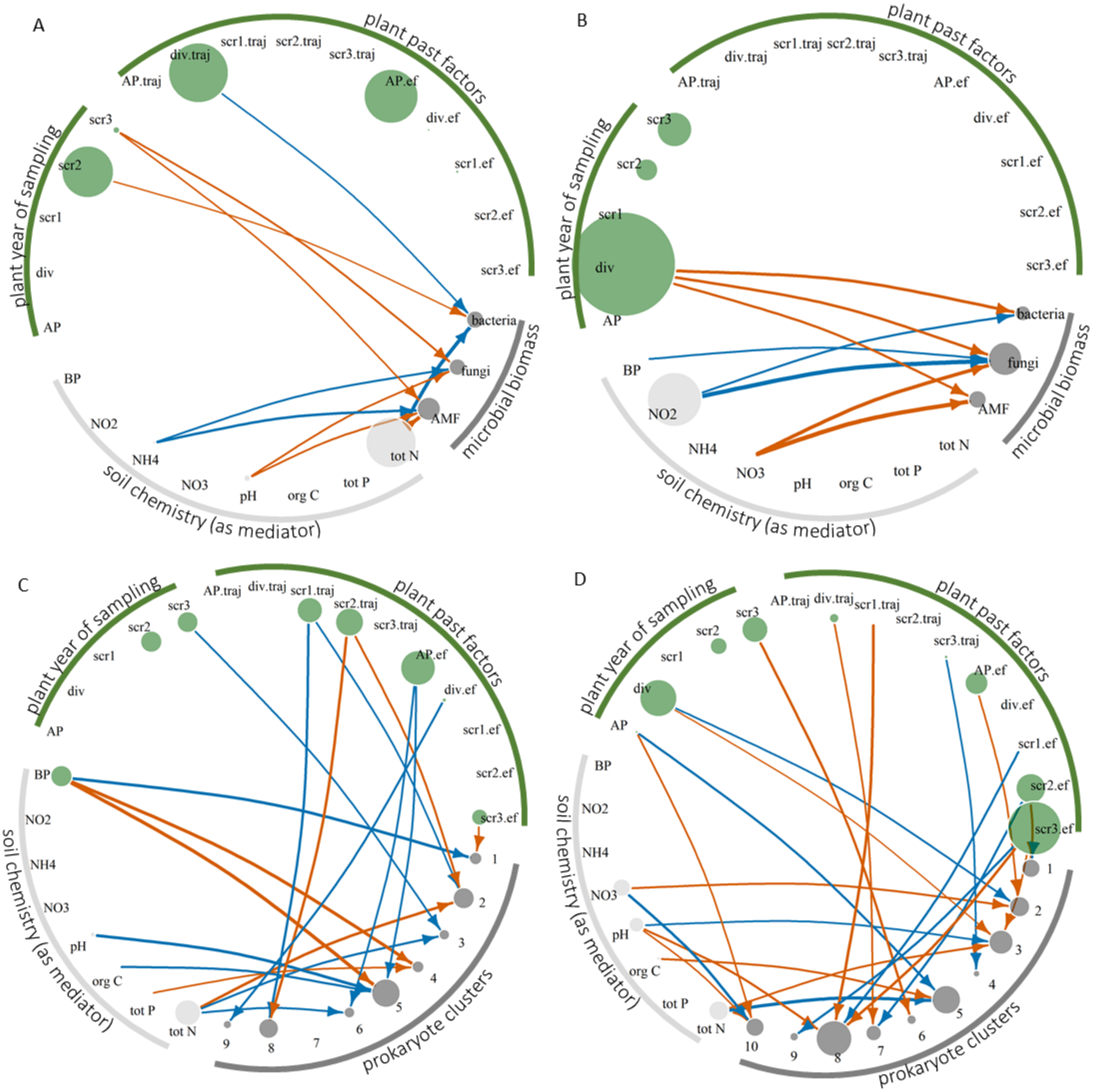


**Figure S11** continued on next page.

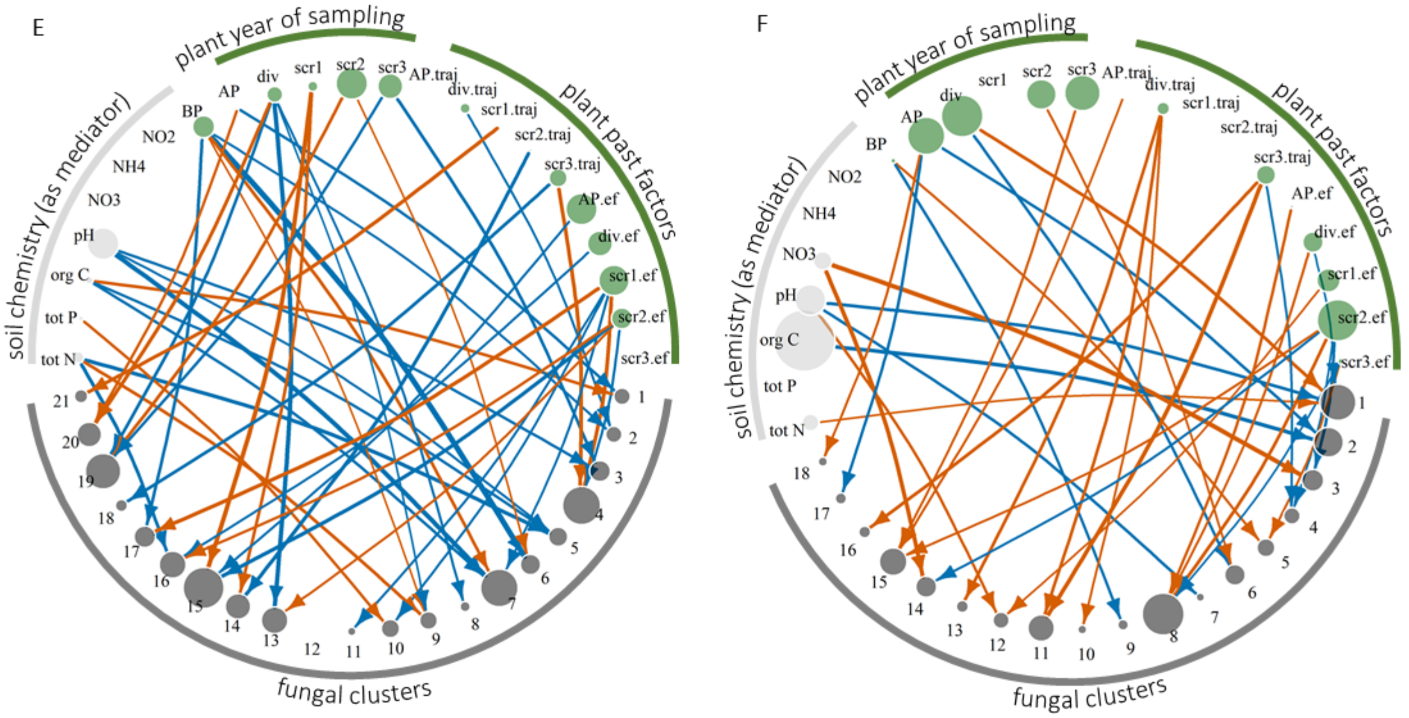


**Figure S11** Significant pathways between plants (green), soil chemistry (light grey) and microbial parameters (dark grey) obtained from structural equation models in (left; A, C, E) natural grassland soil and (right; B, D, F) abandoned arable soil. Incorporated microbial parameters were (A, B) microbial biomass, (B, D) prokaryote network clusters and (E, F) fungal network clusters. Negative pathways are represented in vermillion, positive in blue. Arrows indicate the direction of the pathways and the width of the arrows its effect size. Plant vertice sizes indicate the summed direct and indirect pathway effect sizes onto microbial parameters. Soil chemical vertices indicate only the summed *in*direct pathway effect sizes. Microbial vertices indicate the summed direct and *in*direct pathway effect sizes that these microbial parameters were affected by. All summed pathway effect sizes were scaled to the size of the microbial parameters involved. For plant-soil chemical pathways, see Fig S12.

Plant year of sampling factors: AP – aboveground productivity, div – plant diversity, scr1 – plant composition DCA1 related to species residence period, scr2 – plant composition DCA2 related to species soil resource optima, scr3 – plant composition DCA3 related to community legume cover. Plant past factors: AP.traj – aboveground productivity trajectory, div.traj – plant diversity trajectory, scr1.traj – plant compositional DCA1 trajectory, scr2.traj – plant compositional DCA2 trajectory, scr3.traj – plant compositional DCA3 trajectory, AP.ef – effect size of the start of invasion on aboveground productivity, div.ef - effect size of the start of invasion on plant diversity, scr1.ef - effect size of the start of invasion on DCA1, scr2.ef - effect size of the start of invasion on DCA2, scr3.ef - effect size of the start of invasion on DCA3; soil chemistry: BP – belowground productivity. Only significant pathways included ($p < 0.05$; $n = 30$ communities).

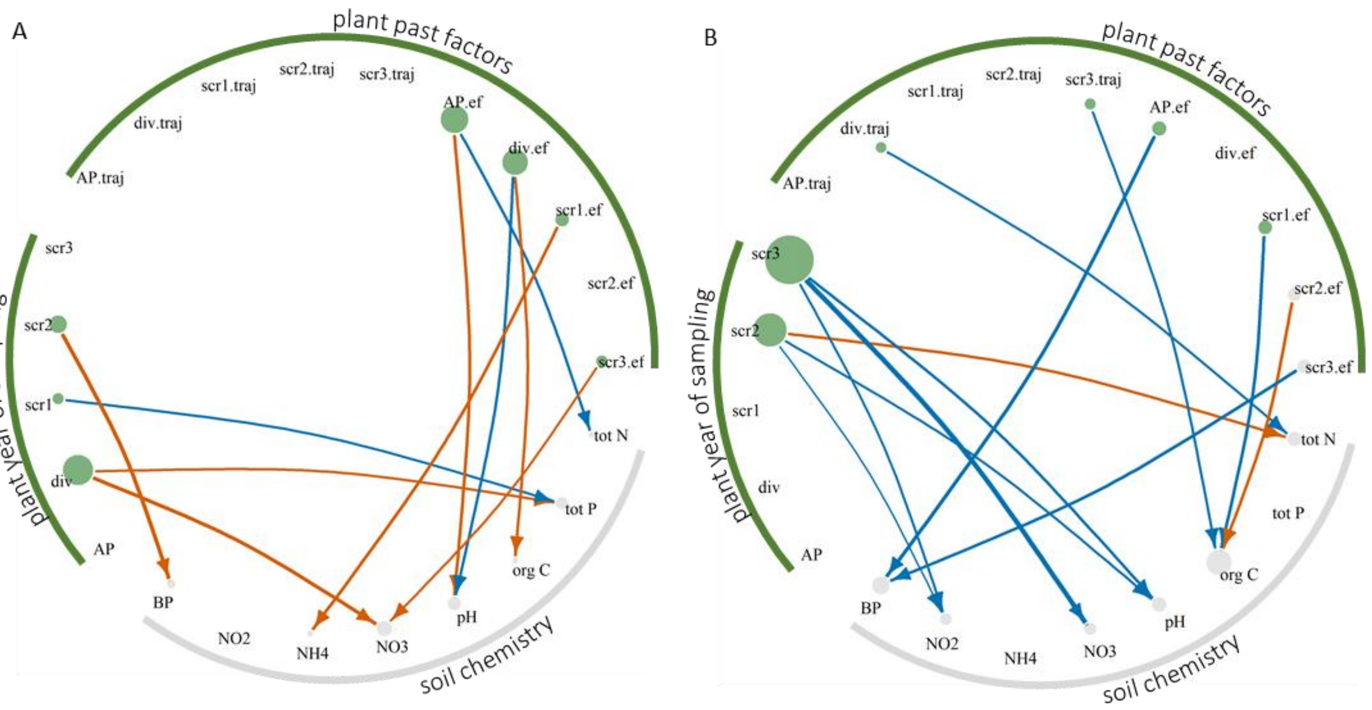


**Figure S12** Significant pathways between plants (green) and soil chemistry (light grey) obtained from structural equation models in (A) natural grassland soil and (B) abandoned arable soil. Negative pathways are represented in vermillion, positive in blue. Arrows indicate the direction of the pathways and the width of the arrows its effect size. Plant vertice sizes indicate the summed pathway effect sizes of the parameter onto all soil chemical properties. Soil chemical vertice sizes indicate the summed pathway effect sizes of all plant community parameters the chemical parameter was affected by.

Plant year of sampling factors: AP – aboveground productivity, div – plant diversity, scr1 – plant composition DCA1 related to species residence period, scr2 – plant composition DCA2 related to species soil resource optima, scr3 – plant composition DCA3 related to community legume cover. Plant past factors: AP.traj – aboveground productivity trajectory, div.traj – plant diversity trajectory, scr1.traj – plant compositional DCA1 trajectory, scr2.traj – plant compositional DCA2 trajectory, scr3.traj – plant compositional DCA3 trajectory, AP.ef – effect size of the start of invasion on aboveground productivity, div.ef - effect size of the start of invasion on plant diversity, scr1.ef - effect size of the start of invasion on DCA1, scr2.ef - effect size of the start of invasion on DCA2, scr3.ef - effect size of the start of invasion on DCA3; soil chemistry: BP – belowground productivity. Only significant pathways included ($p < 0.05$; $n = 30$ communities).

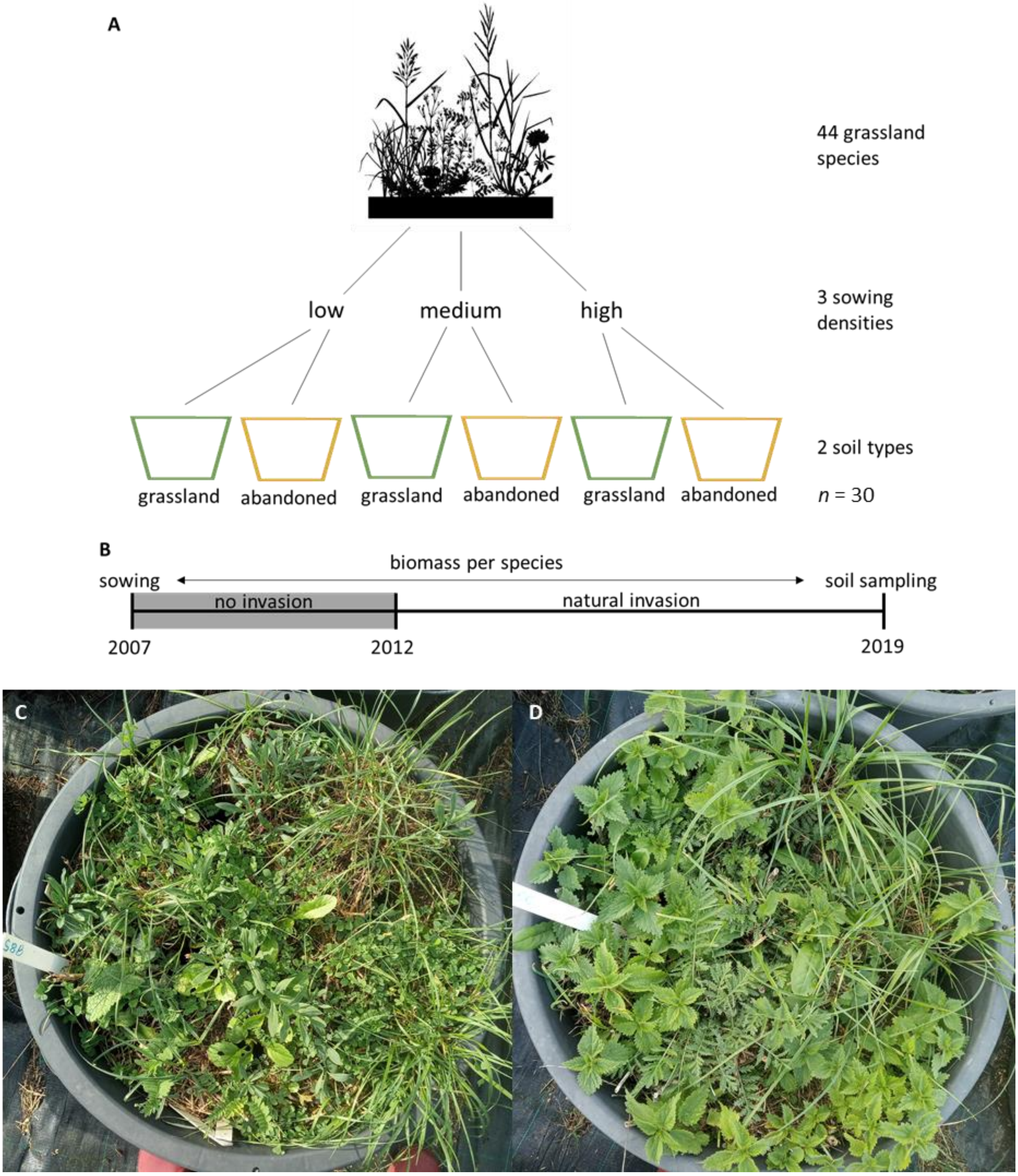


**Figure S13** (A) Experimental design, (B) timeline of the experiment and two typical images of a (C) natural grassland and (D) abandoned arable plant communities (summer 2020). In A, sowing densities represent 25% (low), 100% (medium) and 400% (high) of the natural seed densities as determined in a natural grassland community were the 44 plant species naturally coexist. Grassland soil was taken from a natural grassland and abandoned soil from an arable field that was abandoned in the 1950s (see also Münzbergová, 2012). In B, plant communities were sown in 2007 followed by a 5 year period in which invasion of other plant species was avoided. From 2012 to 2019, natural invasion occurred. In 2019 after the growing season, soil cores for chemical and microbial analysis were taken. For sown and invaded plant species, see Table S9.

**Supplementary tables**

**Table S1** Microbial co-occurrence network parameters typically associated with network stability for networks based on 16S and ITS amplicon sequencing

| | **Prokaryote network** | | **Fungal network** | |
|---|---|---|---|---|
| | Natural grassland | Abandoned arable | Natural grassland | Abandoned arable |
| Number of nodes | 1008 | 1024 | 403 | 455 |
| Number of edges | 10209 | 10079 | 1470 | 1874 |
| Average number of edges per node | 20.3 | 19.7 | 7.3 | 8.2 |
| Negative edges | 46% | 46% | 36% | 33% |
| Edge betweenness | 132 | 139 | 467 | 510 |
| Average edge weight | 0.07 | 0.07 | 0.08 | 0.08 |
| Number of clusters | 9 | 10 | 21 | 18 |
| Clustering coefficient | 0.36 | 0.35 | 0.54 | 0.56 |

**Table S2** Dominant phyla, orders and families in soil prokaryote clusters of plant communities grown on natural grassland soil.

| Cluster | Dominant phyla (>20%) | Dominant orders (>9%) | Dominant families (>9%) | Relates to | Putative metabolic traits and functions |
|---|---|---|---|---|---|
| **1** (95 OTUs) | *Proteobacteria (77.6%)* | *Sphingomondales (30.4%)*<br>*Pseudomonadales (18.6%)*<br>*Xanthomonadales (13.1%)* | *Sphingomonadaceae (30.3%)*<br>*Pseudomonadaceae (18.6%)*<br>*Xanthomonadaceae (13.1%)* | BP (+), invasion effect size DCA3 (-) | Abundant, ubiquitous soil and plant rhizosphere chemoheterotrophs; metabolically diverse, fast growing, various plant-phytopathogens and putative growth promoters[1–3]. Enriched in natural grassland soil. |
| **2** (263 OTUs) | *Actinobacteria (57.7%)*<br>*Proteobacteria (32.6%)* | *Micrococcales (16.2%)*<br>*Solibrubacterales (12.2%)*<br>*Rhizobiales (11.1%)* | *Micrococcaceae (10.0%)*<br>*Solirubrobacteraceae (9.6%)* | Total N (-), DCA1 trajectory (+), DCA2 trajectory (-) | Diverse group of chemoheterotrophs, various involved in N-fixation[4,5]. Many unknown, enriched in natural grassland soil. |
| **3** (244 OTUs) | *Bacteroidetes (44.6%)*<br>*Proteobacteria (26.9%)* | *Chitinophagales (24.9%)*<br>*Cytophagales (17.3%)* | *Chitinophagaceae (24.9%)*<br>*Fulvivirgaceae (16.3%)* | Total N (+), DCA3 (+) | Diverse group of generalist, chemoorganotrophs, degrading organic material, amongst others chitin[6,7]. Present in both soils. |
| **4** (38 OTUs) | *Proteobacteria (80.6%)* | *Burkholderiales (38.0%)*<br>*Nitrosomonadales (13.3%)* | *Burkholderiaceae (33.9%)*<br>*Nitrosomonadaceae (13.3%)* | PO4 (-), BP (-) | Metabolically extremely diverse group including amongst others lithoautotrophic ammonia oxidizers[8,9]. Present in both soils. |
| **5** (61 OTUs) | *Proteobacteria (66.1%)* | *Rhizobiales (27.6%)*<br>*Nevskiales (21.3%)* | *Steroidobacteraceae (21.3%)*<br>*Hyphomicrobiaceae (14.7%)* | Organic C (+), pH (+), BP (-), invasion effect size AP (+) | N-fixing taxa and taxa with unclear function[4]. Present in both soils. |
| **6** (5 OTUs) | *Thaumarchaeota (98.7%)* | *Nitrososphaerales (98.7%)* | *Nitrososphaeraceae (98.7%)* | Total N (+), invasion effect size AP (+) | Ammonia oxidising archaea[10]. Enriched in natural grassland soil. |
| **7** (120 OTUs) | *Proteobacteria (57.3%)* | *All below 9%* | *All below 9%* | Nothing | Unclear specialists. Present in both soils. |
| **8** (2 OTUs) | *Actinobacteria (100%)* | *Acidimicrobiales (60.5%)*<br>*Streptomycetales (39.5%)* | *Acidimicrobiaceae (60.5%)*<br>*Streptomycetaceae (39.5%)* | DCA1 trajectory (+), DCA2 trajectory (-) | Specialists. Obligate acidophilic, reducing ferric iron (*Aciditerrimonas*) and common root endophytes producing mycelium and antibiotic secondary metabolites (*Streptomyces*)[11,12]. Present in both soils. |
| **9** (2 OTUs) | *Actinobacteria (52.3%)*<br>*Proteobacteria (47.7%)* | *Thermoleophilales (52.3%)*<br>*Rhodobacterales (47.7%)* | *Thermoleophilaceae (52.3%)*<br>*Rhodobacteraceae (47.7%)* | Invasion effect size diversity (+) | Globally distributed specialists with unclear function[13]. Present in both soils. |

Percentages indicate the relative read abundance of the phylogenetic group within the cluster. For cluster specialisation and enrichment metrices, see Fig S8-S9.

**Table S3** Dominant phyla, orders and families in soil prokaryote clusters of plant communities grown on abandoned arable soil.

| Cluster | Dominant phyla (>20%) | Dominant orders (>10%) | Dominant families (>9%) | Relates to | Putative metabolic traits and function |
|---|---|---|---|---|---|
| **1** (232 OTUs) | *Proteobacteria (50.4%)*<br>*Actinobacteria (20.1%)* | *Spingomonadales (18.2%)* | *Sphingomonadaceae (16.8%)* | Invasion effect size DCA1 (-), invasion effect size DCA2 (+) | Diverse group of generalist, ubiquitous soil and plant rhizosphere chemoheterotrophs; metabolically diverse and putative growth promoters[3]. Many unknown, enriched in abandoned soil. |
| 2 (226 OTUs) | *Proteobacteria (54.2%)* | *Burkholderiales (14.0%)*<br>*Chitinophagales (13.1%)* | *Chitinophagaceae (13.1%)* | NO3 (-), plant diversity (+), invasion effect size AP (-) | Diverse group of bacteria including chitin-degraders[6]. Many unknown, enriched in abandoned soil. |
| **3** (245 OTUs) | *Proteobacteria (28.5%)*<br>*Actinobacteria (21.4%)* | *Nitrososphaerales (13.6%)*<br>*Rhizobiales (12.8%)*<br>*Cytophagales (10.6%)* | *Nitrososphaeraceae (13.6%)*<br>*Fulvivirgaceae (10.4%)*<br>*Hyphomicrobiaceae (9.9%)* | Total N (-), pH (+), plant diversity (-), invasion effect size DCA3 (-) | Diverse group of ammonia oxidising archaea, organic matter degraders and N-fixing taxa[4,10]. Enriched in abandoned soil. |
| **4** (25 OTUs) | *Actinobacteria (44.2%)*<br>*Acidobacteria (25.7%)*<br>*Proteobacteria (22.2%)* | *Solirubrobacterales (15.8%)*<br>*Xanthomonadales (13.2%)*<br>*Gaiellales (12.5%)* | *Solirubrobacteraceae (15.8%)*<br>*Xanthomonadaceae (13.2%)*<br>*Gaiellaceae (12.5%)* | DCA3 trajectory (+) | Generalist, chemoheterotrophs, various plant-phytopathogens[2,14]. Enriched in abandoned soil. |
| **5** (116 OTUs) | *Proteobacteria (52.0%)*<br>*Actinobacteria (22.2%)* | *Solirubrobacterales (12.4%)*<br>*Sphingomonadales (12.4%)*<br>*Rhizobiales (11.1%)* | *Solirubrobacteraceae (12.4%)*<br>*Sphingomonadaceae (12.0%)*<br>*Chitinophagaceae (9.0%)* | Total N (+), organic C (-), AP (+) | Specialist, ubiquitous soil and plant rhizosphere chemoheterotrophs including chitin-degraders; metabolically diverse and putative growth promoters[3]. Enriched in natural grassland soil. |
| **6** (11 OTUs) | *Proteobacteria (60.5%)* | *Gemmatimonadales (17.3%)*<br>*Myxococcales (16.6%)*<br>*Burkholderiales (14.0%)*<br>*Micropepsales (12.8%)*<br>*Desulfuromonadales (11.9%)*<br>*Chthoniobacterales (11.0%)* | *Gemmatimonadaceae (17.3%)*<br>*Comamonadaceae (14.0%)*<br>*Micropepsaceae (12.8%)*<br>*Chthoniobacteraceae (11.0%)* | DCA3 (-) | Specialist, ubiquitous soil bacteria; metabolically diverse, including organic compound degraders and nitrate reducing bacteria[15–19]. Present in both soils. |
| **7** (4 OTUs) | *Proteobacteria (45.9%)*<br>*Acidobacteria (27.4%)* | *Rhizobiales (45.9%)*<br>*Verrucomicrobiales (18.4%)* | *Methylocystaceae (45.9%)*<br>*Verrucomicrobia subdivision 3 (18.4%)* | Plant diversity trajectory (-), DCA2 trajectory (+) | Ubiquitous soil bacteria, including type II methanotrophs (aerobic methane oxidisers)[20,21]. Present in both soils. |
| **8** (2 OTUs) | *Bacteroidetes (63.6%)*<br>*Acidobacteria (36.4%)* | *Chitinophagales (63.6%)* | *Chitinophagaceae (63.6%)* | pH (-), DCA1 trajectory (-), invasion effect size DCA1 (+), invasion effect size DCA3 (-) | Specialist, chitin-degraders[6]. Present in both soils. |
| **9** (2 OTUs) | *Acidobacteria (76.8%)* | *Burkholderiales (14.8%)* | *Alcaligenaceae (14.8%)* | Invasion effect size DCA3 (+) | Unclear and many unknown specialists. Enriched in abandoned soil. |
| **10** (1 OTU) | *Actinobacteria (68.0%)*<br>*Bacteroidetes (32.0%)* | *Acidimicrobiales (68.0%)*<br>*Chitinophagales (32.0%)* | *Acidimicrobiaceae (68.0%)*<br>*Chitinophagaceae (32.0%)* | pH (-), NO3 (+), AP (-) | Specialist, obligate acidophilic, ferric iron reducing bacteria (*Aciditerrimonas*) and chitin-degraders[6,12]. Enriched in natural grassland soil. |

Percentages indicate the relative read abundance of the phylogenetic group within the cluster. For cluster specialisation and enrichment metrices, see Fig S8-S9.

**References Tables S2-S3**

**Table S4** Dominant phyla, orders, families and putative traits in soil fungal clusters of plant communities grown on natural grassland soil.

| Cluster | Dominant phyla (>20%) | Dominant orders (>9%) | Dominant families (>9%) | Relates to | Dominant fungal traits (>9%) | Putative metabolic traits and functions |
|---|---|---|---|---|---|---|
| **1** (6 OTUs) | *Ascomycota (41.0%)* *Mucoromycota (37.5%)* | *Mucorales (37.5%)* *Archaeorhizomycetales (15.7%)* *Mortierellales (15.0%)* | *Mucoraceae (37.5%)* *Archaeorhizomycetaceae (15.7%)* *Mortierellaceae (15.0%)* | Organic C (-), AP (+) | Soil saprotroph (68.2%) | Soil saprotrophs. Present in both soils. |
| **2** (4 OTUs) | *Ascomycota (89.0%)* | *Filobasidiales (9.4%)* | *Piskurozymaceae (9.4%)* | BP (+), plant diversity trajectory (+) | Soil saprotroph (9.4%) | Abundant, generalist and largely unknown *Ascomycota*. Likely plant root associated. Enriched in natural grassland soil. |
| **3** (46 OTUs) | *Mortierellomycota (50.7%)* *Ascomycota (36.3%)* | *Mortierellales (50.7%)* | *Mortierellaceae (50.7%)* | pH (+), DCA3 (+) | Soil saprotroph (52.7%) Wood saprotroph (10.9%) | Diverse group of generalist saprotrophs. Enriched in natural grassland soil. |
| **4** (7 OTUs) | *Ascomycota (77.3%)* *Basidiomycota (22.7%)* | *Archaeorhizomycetales (54.7%)* *Agaricales (21.9%)* *Geoglossales (18.1%)* | *Archaeorhizomycetaceae (54.6%)* *Hygrophoraceae (19.2%)* *Geoglossaceae (18.1%)* | DCA3 trajectory (-), invasion effect size DCA1 (-), invasion effect size DCA2 (+) | Soil saprotroph (94.7%) | Abundant soil saprotrophs. Enriched in natural grassland soil. |
| **5** (17 OTUs) | *Ascomycota (52.5%)* *Mortierellomycota (42.5%)* | *Mortierellales (42.5%)* *Hypocreales (35.8%)* | *Mortierellaceae (42.5%)* *Nectriaceae (33.4%)* | Total N (+), organic C (+), pH (+) | Plant pathogen (36.5%) Soil saprotroph (35.4%) | Relative diverse and abundant, generalist soil saprotrophs and plant pathogens (*Fusarium, Ilyonectria, Verticillium, Leptosphaeria, Gibberella*). Present in both soils. |
| **6** (13 OTUs) | *Ascomycota (82.2%)* | *Helotiales (57.5%)* *Tremellales (16.5%)* *Capnodiales (12.0%)* *Thelebolales (11.2%)* | *Bulleribasidiaceae (16.5%)* *Cladosporiaceae (12.0%)* *Pseudeurotiaceae (11.2%)* | BP (+), DCA2 (-) | Soil saprotroph (27.9%) Litter saprotroph (15.3%) | Relative abundant, generalist saprotrophs likely plant root associated. Enriched in natural grassland soil. |
| **7** (10 OTUs) | *Ascomycota (86.4%)* | *Pezizales (50.1%)* *Mortierellales (11.4%)* | *Pyronemataceae (46.3%)* *Mortierellaceae (11.4%)* | Organic C (+), pH (+), BP (-), invasion effect size DCA1 (+) | Soil saprotroph (13.6%) Plant pathogen (9.2%) | Soil saprotrophs and plant pathogens (*Fusarium, Lectera*). Present in both soils. |
| **8** (5 OTUs) | *Ascomycota (69.5%)* *Basidiomycota (23.7%)* | *Geoglossales (56.0%)* *Agaricales (23.7%)* | *Geoglossaceae (56.0%)* *Clavariaceae (23.7%)* | Plant diversity (+) | Soil saprotroph (79.7%) | Soil saprotrophs. Present in both soils. |

**Table S4 continued.**

| Cluster | Dominant phyla (>20%) | Dominant orders (>9%) | Dominant families (>9%) | Relates to | Dominant fungal traits | Putative function |
|---|---|---|---|---|---|---|
| **9** (13 OTUs) | *Ascomycota (84.8%)* | *Archaeorhizomycetales (42.3%)* | *Archaeorhizomycetaceae (42.3%)* | Total N (-), BP (-), plant diversity (+) | Soil saprotroph (51.2%) | Soil saprotrophs. Enriched in natural grassland soil. |
| **10** (19 OTUs) | *Ascomycota (55.8%)* | *Pleosporales (38.0%)* *Agaricales (13.5%)* | *Periconiaceae (21.0%)* *Hygrophoraceae (13.0%)* | PO4 (-), invasion effect size DCA1 (+) | Soil saprotroph (22.4%) | Diverse group of unknown fungi and soil saprotrophs. Present in both soils. |
| **11** (11 OTUs) | *Ascomycota (64.1%)* | *Pleosporales (35.9%)* *Helotiales (14.3%)* | *Didymellaceae (21.2%)* *Sclerotiniaceae (13.8%)* *Pleosporaceae (11.2%)* | Invasion effect size DCA1 (+) | Plant pathogen (46.5%) | Plant pathogens (*Phoma*, *Botrytis*, *Stemphylium*, *Ophiosphaerella*). Enriched in natural grassland soil. |
| **12** (5 OTUs) | *Mucoromycota (49.0%)* *Ascomycota (30.2%)* | *Mucorales (49.0%)* *Pleosporales (21.1%)* | *Mucoraceae (49.0%)* *Melanommataceae (21.1%)* | - | Soil saprotroph (50.4%) Plant pathogen (21.1%) | Specialist soil saprotrophs and plant pathogens (*Herpotrichia*). Present in both soils. |
| **13** (8 OTUs) | *Ascomycota (81.0%)* | *Pleosporales (61.2%)* | *Phaeosphaeriaceae (32.2%)* *Pleosporales (24.5%)* | Plant diversity (+), invasion effect size DCA2 (-) | Plant pathogen (37.7%) | Plant pathogens (*Paraphoma*, *Septoria*, *Plenodomus*). Present in both soils. |
| **14** (13 OTUs) | *Ascomycota (82.3%)* | *Pleosporales (33.0%)* *Geoglossales (19.7%)* | *Didymellaceae (25.6%)* *Geoglossaceae (19.7%)* | DCA1 (-), DCA2 trajectory (+) | Soil saprotroph (28.6%) Plant pathogen (26.7%) | Soil saprotrophs and plant pathogens (*Stagonosporopsis*, *Plectosphaerella*). Present in both soils. |
| **15** (8 OTUs) | *Ascomycota (79.9%)* | *Chaetothyriales (41.7%)* *Pleosporales (30.7%)* | *Trichomeriaceae (37.3%)* *Didymellaceae (25.8%)* | DCA1 (-), invasion effect size AP (+), invasion effect size DCA2 (+) | Unspecified saprotroph (37.3%) Plant pathogen (36.3%) | Generalist saprotrophs and plant pathogens (*Ascochyta*, *Gibberella*, *Coniosporium*). Enriched in natural grassland soil. |
| **16** (9 OTUs) | *Ascomycota (69.4%)* | *Pleosporales (44.0%)* | *Unidentified (76.3%)* | Total N (+), invasion effect size diversity (+), invasion effect size DCA2 (-) | Plant pathogen (11.1%) | Possible plant pathogens as order contains various putative plant pathogenic groups; also many unknown. Enriched in natural grassland soil. |

**Table S4 continued.**

| Cluster | Dominant phyla (>20%) | Dominant orders (>9%) | Dominant families (>9%) | Relates to | Dominant fungal traits | Putative function |
|---|---|---|---|---|---|---|
| **17** (9 OTUs) | *Ascomycota (72.1%)* *Basidiomycota (25.9%)* | *Archaeorhizomycetales (58.2%)* *Trechisporales (12.2%)* | *Archaeorhizomycetaceae (58.2%)* | BP (+), invasion effect size DCA1 (-) | Soil saprotroph (62.7%) Wood saprotroph (9.1%) | Saprotrophs likely associated with plant roots. Present in both soils. |
| **18** (10 OTUs) | *Ascomycota (32.3%)* *Basidiomycota (27.9%)* | *Sebacinales (18.5%)* *Mortierallales (14.1%)* *Rhizophlyctidales (9.9%)* | *Sebacinaceae (18.5%)* *Mortierellaceae (14.1%)* *Rhizophlyctidaceae (9.9%)* | DCA3 trajectory (+) | Litter saprotroph (22.8%) Soil saprotroph (14.1%) | Litter and soil saprotrophs. Present in both soils. |
| **19** (13 OTUs) | *Ascomycota (57.4%)* *Basidiomycota (38.0%)* | *Hypocreales (21.0%)* *Agaricales (15.1%)* *Chaetothyriales (13.7%)* *Cantharellales (9.8%)* | *Ceratobasidiaceae (9.8%)* | Plant diversity (+), DCA2 (-), DCA3 (+) | Litter saprotroph (17.1%) Animal parasite (11.0%) Soil saprotroph (10.9%) | Relative diverse group of saprotrophs including putative nematode parasites. Enriched in natural grassland soil. |
| **20** (1 OTU) | *Kickxellomycota (100%)* | *Kickxellales (100%)* | *Kickxellaceae (100%)* | AP (-), plant diversity (-) | Soil saprotroph (100%) | Generalist soil saprotrophs. Enriched in abandoned soil. |
| **21** (2 OTUs) | *Ascomycota (54.9%)* | *Pleosporales (26.5%)* *Verrucariales (16.7%)* *Orbiliales (11.4%)* | *Phaeosphaeriaceae (26.5%)* *Verrucariaceae (16.7%)* *Orbiliaceae (11.4%)* | DCA1 trajectory (-) | Plant pathogen (26.5%) Lichenized (16.7%) Animal parasite (11.4%) | Specialist lichens, nematode parasites (*Arthrobotrys*) and plant pathogens (*Chaetosphaeronema*). Present in both soils. |

Percentages indicate the relative read abundance of the phylogenetic group within the cluster. Dominant fungal traits were obtained from the FungalTrait database (Põlme, S. et al. FungalTraits: a user-friendly traits database of fungi and fungus-like stramenopiles. Fungal Diversity 105, (2020)). Genera identified as putative plant pathogens presented in brackets. For cluster specialisation and enrichment metrices, see Fig S8-S9.

**Table S5** Dominant phyla, orders, families and putative traits in soil fungal clusters of plant communities grown on abandoned arable soil.

| Cluster | Dominant phyla (>20%) | Dominant orders (>9%) | Dominant families (>9%) | Relates to | Dominant fungal traits | Putative function |
|---|---|---|---|---|---|---|
| **1** (56 OTUs) | *Mortierellomycota (46.8%)*<br>*Ascomycota (43.8%)* | *Mortierellales (46.8%)*<br>*Pezizales (18.4%)* | *Mortierellaceae (46.8%)*<br>*Pyronemataceae (16.2%)* | Total N (-), pH (+), plant diversity (-), invasion effect size diversity (+) | Soil saprotroph (52.8%) | Diverse group of generalist soil saprotrophs. Enriched in abandoned soil. |
| **2** (48 OTUs) | *Ascomycota (61.3%)*<br>*Mortierellomycota (34.8%)* | *Mortierellales (34.8%)*<br>*Hypocreales (18.7%)* | *Mortierellaceae (34.8%)*<br>*Nectriaceae (15.8%)* | Organic C (+), AP (+) | Soil saprotroph (38.5%)<br>Plant pathogen (17.8%) | Diverse group of generalist soil saprotrophs and plant pathogens (*Fusarium, Ilyonectria, Nectria, Plenodomus, Thielaviopsis, Lectera, Paraphoma, Ascochyta, Plectosphaerella, Stagonosporopsis*). Enriched in abandoned soil. |
| **3** (27 OTUs) | *Ascomycota (46.3%)*<br>*Basidiomycota (42.3%)* | *Agaricales (35.7%)*<br>*Geoglossales (18.2%)*<br>*Archaeorhizomycetales (12.2%)*<br>*Incertae (9.8%)* | *Clavariaceae (33.0%)*<br>*Geoglossaceae (18.2%)*<br>*Archaeorhizomycetaceae (12.2%)* | NO3 (-), invasion effect size DCA2 (+) | Soil saprotroph (61.4%) | Diverse group of soil saprotrophs. Enriched in abandoned soil. |
| **4** (5 OTUs) | *Ascomycota (62.4%)*<br>*Basidiomycota (37.6%)* | *Agaricales (37.6%)*<br>*Geoglossales (20.5%)*<br>*Eurotiales (14.2%)*<br>*Hypocreales (12.0%)*<br>*Saccharomycetales (11.6%)* | *Tricholomataceae (37.6%)*<br>*Geoglossaceae (20.5%)*<br>*Aspergillaceae (14.2%)*<br>*Nectriaceae (12.0%)*<br>*Debaryomycetaceae (11.6%)* | DCA3 trajectory (+), invasion effect size DCA2 (+) | Litter saprotroph (37.6%)<br>Soil saprotroph (20.5%)<br>Plant pathogen (16.1%)<br>Unspecified saprotroph (14.2%)<br>Nectar/tap saprotroph (11.6%) | Saprotrophs and plant pathogens (*Fusarium, Protomyces*). Present in both soils. |
| **5** (17 OTUs) | *Ascomycota (48.9%)*<br>*Mortierellomycota (23.0%)* | *Hypocreales (34.1%)*<br>*Mortierellales (23.0%)*<br>*Glomerales (9.4%)* | *Mortierellaceae (23.0%)*<br>*Hypocreaceae (18.3%)*<br>*Nectriaceae (15.8%)* | BP (-), invasion effect size DCA3 (-) | Soil saprotroph (23.0%)<br>Mycoparasite (18.3%) | Diverse group of soil saprotrophs and mycoparasites. Present in both soils. |

**Table S5 continued.**

| Cluster | Dominant phyla (>20%) | Dominant orders (>9%) | Dominant families (>9%) | Relates to | Dominant fungal traits | Putative function |
|---|---|---|---|---|---|---|
| **6** (39 OTUs) | *Ascomycota (56.3%)*<br>*Basidiomycota (21.3%)* | *Agaricales (17.4%)*<br>*Sordariales (15.0%)*<br>*Pleosporales (14.2%)*<br>*Orbiliales (12.4%)* | *Agaricaceae (17.1%)*<br>*Orbiliaceae (12.4%)* | Plant diversity (+), DCA2 (-) | Soil saprotroph (29.8%) | Diverse group of soil saprotrophs and small proportion, but high diversity of plant pathogens (*Herpotrichia, Chaetosphaeronema, Verticillium, Stemphylium, Periconia, Leptosphaeria, Fusarium, Alternaria, Gibellulopsis, Botrytis, Gibberella*). Enriched in abandoned soil. |
| **7** (12 OTUs) | *Ascomycota (48.3%)*<br>*Basidiomycota (36.5%)* | *Sebacinales (21.9%)*<br>*Pezizales (19.5%)*<br>*Pleosporales (12.7%)* | *Sebacinaceae (21.9%)*<br>*Pyronemataceae (14.3%)* | pH (+) | Soil saprotroph (25.5%)<br>Litter saprotroph (11.6%) | Soil and litter saprotrophs. Present in both soils. |
| **8** (2 OTUs) | *Kickxellomycota (57.4%)* | *Kickxellales (57.4%)*<br>*Mortierellales (11.9%)* | *Kickxellaceae (57.4%)*<br>*Mortierellaceae (11.9%)* | Invasion effect size AP (-), invasion effect size diversity (-), invasion effect size DCA2 (-), invasion effect size DCA3 (+) | Soil saprotroph (69.3%) | Generalist soil saprotrophs. Present in both soils. |
| **9** (10 OTUs) | *Basidiomycota (42.1%)*<br>*Ascomycota (36.2%)* | *Sebacinales (29.1%)* | *Serendipitaceae (29.1%)* | BP (+) | Root endophyte (29.1%)<br>Litter saprotroph (10.9%) | Generalist root endophytes (*Serendipita*) and litter saprotrophs. Present in both soils. |
| **10** (10 OTUs) | *Ascomycota (88.9%)* | *Pleosporales (39.8%)*<br>*Geoglossales (18.6%)*<br>*Pezizales (9.2%)* | *Didymellaceae (24.1%)*<br>*Geoglossaceae (18.6%)*<br>*Lentitheciaceae (9.2%)*<br>*Pyronemataceae (9.2%)* | Plant diversity trajectory (-) | Soil saprotroph (28.1%)<br>Plant pathogen (24.1%)<br>Root endophyte (9.2%) | Soil saprotrophs, plant pathogens (*Ascochyta*) and root endophytes (*Darksidea*). Present in both soils. |
| **11** (2 OTUs) | *Chytridiomycota (100%)* | *Rhizophydiales (100%)* | *Rhizophydiaceae (100%)* | Plant diversity trajectory (-), DCA3 trajectory (-) | Algal parasite (100%) | Generalist algal parasites. Enriched in natural grassland soil. |
| **12** (8 OTUs) | *Ascomycota (51.2%)*<br>*Olpidiomycota (36.3%)* | *Olpidiales (36.3%)* | *Olpidiaceae (36.3%)* | pH (-), DCA1 trajectory (-) | Algal parasite (36.3%)<br>Litter saprotroph (10.8%) | Specialist litter saprotrophs and algal parasites. Present in both soils. |

**Table S5 continued**

| Cluster | Dominant phyla (>20%) | Dominant orders (>9%) | Dominant families (>9%) | Relates to | Dominant fungal traits | Putative function |
|---|---|---|---|---|---|---|
| **13** (10 OTUs) | *Ascomycota (43.4%)* *Basidiomycota (35.3%)* | *Agaricales (18.9%)* *Mortierellales (15.3%)* *Cantharellales (15.3%)* *Pezizales (14.8%)* *Hypocreales (10.4%)* | *Mortierellaceae (15.3%)* *Cantharellales (15.3%)* *Pyronemataceae (14.8%)* *Marasmiaceae (12.8%)* | Plant diversity trajectory (-) | Soil saprotroph (17.2%) Litter saprotroph (17.1%) Lichen parasite (15.3%) Wood saprotroph (14.3%) | Soil, litter and wood saprotrophs as well as lichen parasites. Present in both soils. |
| **14** (6 OTUs) | *Ascomycota (50.4%)* *Basidiomycota (37.7%)* | *Archaeorhizomycetales (44.6%)* *Agaricales (37.7%)* *Rhizophydiales (10.9%)* | *Archaeorhizomycetaceae (44.6%)* *Clavariaceae (30.5%)* | NO3 (-), invasion effect size DCA2 (+) | Soil saprotroph (82.3%) | Specialist soil saprotrophs. Present in both soils. |
| **15** (6 OTUs) | *Ascomycota (74.9%)* | *Chaetothyriales (19.9%)* *Pleosporales (16.5%)* *Helotiales (15.7%)* *Orbiliales (10.8%)* *Cystofilobasidiales (10.3%)* | *Sporormiaceae (16.5%)* *Orbiliaceae (10.8%))* *Mrakiaceae (10.3%)* | DCA3 (-), AP trajectory (-), invasion effect size DCA2 (-) | Dung saprotroph (16.5%) Plant pathogen (10.3%) Litter saprotroph (9.9%) | Litter and dung saprotrophs, and plant pathogens (*Itersonilia*). Present in both soils. |
| **16** (2 OTUs) | *Basidiomycota (100%)* | *Unidentified (100%)* | *Unidentified (100%)* | DCA3 trajectory (-) | Unknown (100%) | Unknown generalists. Enriched in abandoned soil. |
| **17** (1 OTU) | *Ascomycota (100%)* | *Melanosporales (60.6%)* | *Melanosporaceae (60.6%)* | AP (+) | Mycoparasite (60.6%) | Specialist mycoparasites. Present in both soils. |
| **18** (1 OTU) | *Basidiomycota (53.3%)* *Ascomycota (46.7%)* | *Sebacinales (53.3%)* *Helotiales (46.7%)* | *Serendipitaceae (53.3%)* | AP (-) | Root endophyte (53.3%) | Specialist root endophytes (*Serendipita*). Enriched in abandoned soil. |

Percentages indicate the relative read abundance of the phylogenetic group within the cluster. Dominant fungal traits were obtained from the FungalTrait database (Põlme, S. et al. FungalTraits: a user-friendly traits database of fungi and fungus-like stramenopiles. Fungal Diversity 105, (2020)). Genera identified as putative plant pathogens presented in brackets. For cluster specialisation and enrichment metrices, see Fig S8-S9.

**Table S6** Marginal $R^2$ of soil chemistry, microbial biomass and clusters from SEM models in natural grassland and abandoned arable plant communities.

| | Natural grassland | Abandoned arable | | Natural grassland | Abandoned arable |
|---|---|---|---|---|---|
| **Soil chemistry** | | | **Fungal clusters** | | |
| BP | 0.31 | 0.43 | 1 | 0.34 | **0.59** |
| Total N | 0.18 | 0.24 | 2 | **0.29** | **0.41** |
| Total P | 0.21 | 0.00 | 3 | **0.34** | **0.38** |
| Organic C | 0.16 | 0.50 | 4 | **0.44** | 0.20 |
| NO3 | 0.43 | 0.47 | 5 | **0.41** | 0.34 |
| NH4 | 0.21 | 0.00 | 6 | **0.61** | 0.28 |
| NO2 | 0.00 | 0.19 | 7 | 0.53 | 0.15 |
| pH | 0.26 | 0.25 | 8 | 0.15 | 0.38 |
| **PLFA/NLFA** | | | 9 | 0.61 | 0.18 |
| Bacterial biomass | 0.52 | 0.49 | 10 | 0.31 | 0.12 |
| Fungal biomass | 0.41 | 0.54 | 11 | 0.10 | 0.28 |
| AMF biomass | 0.41 | 0.48 | 12 | 0.00 | 0.20 |
| **Prokaryote clusters** | | | 13 | 0.35 | 0.22 |
| 1 | **0.30** | **0.35** | 14 | 0.33 | 0.36 |
| 2 | **0.41** | **0.54** | 15 | 0.62 | 0.39 |
| 3 | **0.29** | **0.64** | 16 | 0.36 | 0.20 |
| 4 | 0.36 | 0.12 | 17 | 0.33 | 0.18 |
| 5 | 0.35 | 0.4 | 18 | 0.21 | 0.13 |
| 6 | 0.4 | 0.22 | 19 | 0.29 | |
| 7 | 0.00 | 0.29 | 20 | 0.15 | |
| 8 | 0.23 | 0.53 | 21 | 0.21 | |
| 9 | 0.17 | 0.16 | | | |
| 10 | | 0.35 | | | |

$R^2$ in bold belong to large clusters (Fig 3). Note: sowing density was incorporated as a random effect, but in most cases explained no variation. Marginal $R^2$ indicates variation explained by fixed factors only.

**Table S7** Summarised putative effects of the strongest pathways of the plant community in the year of sampling and past on microbial communities in natural grassland soil.

| Time point | Plant parameter | Pathway | Overall microbial parameters | Prokaryote clusters | Fungal clusters | Putative metabolic traits and functions |
|---|---|---|---|---|---|---|
| Year of sampling | AP | direct | - | - | ↑1 ↓20 | ↑ Soil saprotrophs<br>↓ Generalist soil saprotrophs |
| **Year of sampling** | **Plant diversity** | **direct** | ↑fungal diversity | - | ↑8 ↑9<br>↑13 ↑19<br>↓20 | ↑ Metabolically diverse group of bacteria including ammonia oxidizers; diverse group of soil saprotrophs, various plant pathogens, nematode parasites and diverse group unknown fungi<br>↓ Generalist soil saprotrophs |
| | | via ↓P | ↑fungal diversity | ↑4 | ↑10 | |
| Year of sampling | Composition (DCA1) – temporal turnover | direct | ↓prokaryote SI | - | ↓14 ↓15 | ↓ Metabolically diverse group of bacteria including ammonia oxidizers; diverse group of unknown fungi, soil saprotrophs and plant pathogens; generalist saprotrophs and plant pathogens |
| | | via ↑P | ↓fungal diversity | ↓4 | ↓10 | |
| **Year of sampling** | **Composition (DCA2) – soil resource optima** | direct | ↓bacterial biomass<br>↓fungal diversity | - | **↓6** ↓19 | ↑ Metabolically diverse group of bacteria including ammonia oxidizers and N-fixers; soil saprotrophs and plant pathogens<br>↓ Bacterial biomass; ubiquitous soil and plant rhizosphere chemoheterotrophs; abundant, generalist saprotrophs and unknown fungi (likely root-associated); diverse group of soil saprotrophs |
| | | **via ↓BP** | ↑prokaryote diversity | **↓1** ↑4 ↑5 | **↓2** **↓6** ↑7<br>↑9 ↓17 | |
| **Year of sampling** | **Composition (DCA3) – legume cover** | **direct** | ↓fungal biomass<br>↓AMF biomass | **↑3** | **↑3** ↑19 | ↑ Generalist, chemoorganotrophs including chitin degrading bacteria; diverse group of generalist saprotrophs<br>↓ Fungal and AMF biomass |
| Past | AP trajectory | direct | - | - | - | - |
| Past | Plant diversity trajectory | direct | ↑bacterial biomass | - | - | ↑ Bacterial biomass. |
| **Past** | **2012 invasion effect size AP** | direct | ↑fungal diversity | ↑5 ↑6 | ↑15 | ↑ Bacterial, fungal and AMF biomass; ammonia oxidising archaea; N-fixing and diverse group of generalist organic compound degrading bacteria; diverse and abundant, generalist soil saprotrophs and plant pathogens.<br>↓ AMF biomass; N-fixing and diverse group of chemoheterotrophic bacteria; diverse and abundant, generalist soil saprotrophs and plant pathogens. |
| | | **via ↑N** | ↑bacterial biomass<br>↓AMF biomass<br>↑prokaryote SI | **↓2** **↑3** ↑6 | **↑5** ↓9<br>↑16 | |
| | | **via ↓pH** | ↑fungal biomass<br>↑AMF biomass | ↓5 | **↓3** **↓5** ↓7 | |
| **Past** | **2012 invasion effect size diversity** | direct | ↓fungal SI | ↑9 | ↑16 | ↑ N-fixing and ubiquitous specialist bacteria; diverse group of soil saprotrophs and plant pathogens<br>↓ Fungal and AMF biomass; N-fixing bacteria; some soil saprotrophs and plant pathogens |
| | | **via ↓org C** | ↑prokaryote SI | ↓5 | ↑1 **↓5** ↓7 | |
| | | **via ↑pH** | ↓fungal biomass<br>↓AMF biomass | ↑5 | **↑3** **↑5** ↑7 | |

Pathways in bold indicate pathways with a relative contribution > 5% (see Fig 6). Clusters in bold indicate dominant clusters (see Fig 3). Clusters with a thick underline were classified as relative habitat generalists and clusters with a wave underline were classified as relative habitat specialists (see Fig S9). Clusters highlighted in green were enriched in natural grassland soil and clusters highlighted in red were enriched in abandoned arable soil (see Fig S10). For cluster specific information, see Table S2 and S4.

**Table S7 continued.**

| Time point | Plant parameter | Pathway | Overall microbial parameters | Prokaryote clusters | Fungal clusters | Putative metabolic traits and functions |
|---|---|---|---|---|---|---|
| Past | **DCA1 trajectory** | **direct** | - | **↑2** ↑8 | ↓21 | ↑ Diverse group of chemoheterotrophs, various involved in N-fixation; specialist obligate acidophiles and root endophytes<br>↓ Specialist lichens, nematode parasites and plant pathogens |
| Past | **DCA2 trajectory** | **direct** | - | **↓2** ↓8 | ↑14 | ↑ Soil saprotrophs and plant pathogens<br>↓ Diverse group of chemoheterotrophs, various involved in N-fixation; specialist obligate acidophiles and root endophytes |
| Past | **DCA3 trajectory** | **direct** | - | - | **↓4** ↑18 | ↑ Litter and soil saprotrophs.<br>↓ Abundant soil saprotrophs. |
| Past | **2012 invasion effect size DCA1** | **direct** | - | - | **↓4** ↑7 ↑10 ↑11 ↓17 | ↑ Diverse group of unknown fungi, soil saprotrophs and plant pathogens.<br>↓ Fungal and AMF biomass; abundant soil saprotrophs. |
| | | via ↓NH4 | ↓fungal biomass ↓AMF biomass | - | - | |
| Past | **2012 invasion effect size DCA2** | **direct** | - | - | **↑4** ↓13 ↑15 ↓16 | ↑ Abundant soil saprotrophs, generalist saprotrophs and plant pathogens.<br>↓ Soil saprotrophs and plant pathogens. |
| Past | **2012 invasion effect size DCA3** | **direct** | - | **↓1** | - | ↓ Abundant ubiquitous soil and plant rhizosphere chemoheterotrophs |

Pathways in bold indicate pathways with a relative contribution > 5% (see Fig 6). Clusters in bold indicate dominant clusters (see Fig 3). Clusters with a thick underline were classified as relative habitat generalists and clusters with a wave underline were classified as relative habitat specialists (see Fig S9). Clusters highlighted in green were enriched in natural grassland soil and clusters highlighted in red were enriched in abandoned arable soil (see Fig S10). For cluster specific information, see Table S2 and S4.

**Table S8** Summarised putative effects of the strongest pathways of the plant community in the year of sampling and past on microbial communities in abandoned arable soil.

| Time point | Plant parameter | Pathway | Overall microbial parameters | Prokaryote clusters | Fungal clusters | Putative metabolic traits and functions |
|---|---|---|---|---|---|---|
| Year of sampling | **Aboveground productivity** | **direct** | - | ↑5 ↓10 | **↑2** ↑17<br>↓18 | ↑ Specialist, ubiquitous soil and plant rhizosphere chemoheterotrophs including chitin-degraders; diverse group of generalist soil saprotrophs and plant pathogens; specialist mycoparasites<br>↓ Specialist, obligate acidophiles and chitin-degraders; specialist root endophytes |
| Year of sampling | **Plant diversity** | **direct** | ↓bacterial biomass<br>↓fungal biomass<br>↓AMF biomass | **↑2 ↓3** | **↓1** ↑6 | ↑ Broad group of bacteria including chitin-degraders; diverse group of saprotrophs and plant pathogens<br>↓ Bacterial, fungal and AMF biomass; diverse group of ammonia oxidising archaea, organic matter degraders and N-fixing taxa; diverse group of generalist soil saprothrophs |
| Year of sampling | Composition (DCA1) – temporal turnover | direct | - | - | - | - |
| Year of sampling | **Composition (DCA2) – soil resource optima** | direct | - | - | ↓6 | ↑ Diverse group of ammonia oxidising archaea, organic matter degraders and N-fixing taxa; diverse group of generalist soil saprotrophs<br>↓ Specialist, ubiquitous soil and plant rhizosphere chemoheterotrophs; specialist chitin-degraders and obligate acidophiles; diverse group of soil saprotrophs and plant pathogens; specialist litter saprotrophs and algal parasites |
| | | **via ↓N** | - | **↑3** ↓5 | **↑1** | |
| | | **via ↑pH** | ↓fungi SI | **↑3** ↓8<br>↓10 | **↑1** ↑7<br>↓12 | |
| Year of sampling | **Composition (DCA3) – legume cover** | direct | ↓AMF biomass | ↓6 | ↓15 | ↑ Diverse group of ammonia oxidising archaea, organic matter degraders and N-fixing taxa; diverse group of generalist saprotrophs<br>↓ Fungal and AMF biomass; abundant, broad group of bacteria including chitin-degraders; specialist, ubiquitous soil bacteria including organic compound and chitin degraders, nitrate reducing and obligate acidophiles; diverse group of soil saprotrophs; specialist saprotrophs and algal parasites; plant pathogens |
| | | **via ↑NO3** | ↓fungal biomass<br>↓AMF biomass | **↓2** | **↓3** ↓14 | |
| | | **via ↑pH** | ↓fungi SI | **↑3** ↓8<br>↓10 | **↑1** ↑7<br>↓12 | |
| Past | AP trajectory | direct | - | - | ↓15 | ↓ litter and dung saprotrophs, and plant pathogens |
| Past | Plant diversity trajectory | direct | - | ↓7 | - | ↑ Specialist, ubiquitous soil and plant rhizosphere chemoheterotrophs.<br>↓ Diverse group of ammonia oxidising archaea, organic matter degraders, N-fixing taxa and methanotrophs; diverse group of generalist soil saprotrophs. |
| | | via ↑N | - | **↓3** ↑5 | **↓1** | |
| Past | **2012 invasion effect size AP** | **direct** | ↓prokaryote SI | **↓2** | **↓8** | ↑ Fungal biomass; generalist root endophytes and litter saprotrophs.<br>↓ Diverse group of bacteria including chitin-degraders; diverse group of soil saprotrophs and mycoparasites; generalists soil saprotrophs. |
| | | via ↑BP | ↑fungal biomass | - | ↓5 ↑9 | |
| | | via ↓NO2 | ↓bacterial biomass<br>↓fungal biomass | - | - | |

Pathways in bold indicate pathways with a relative contribution > 5% (see Fig 6). Clusters in bold indicate dominant clusters (see Fig 3). Clusters with a thick underline were classified as relative habitat generalists and clusters with a wave underline were classified as relative habitat specialists (see Fig S9). Clusters highlighted in green were enriched in natural grassland soil and clusters highlighted in red were enriched in abandoned arable soil (see Fig S10). For cluster specific information, see Table S3 and S5.

**Table S8** Summarised putative effects of the most important plant community parameters in the year of sampling and past effects on microbial soil legacies in abandoned arable soil communities

| Time point | Plant parameter | Pathway | Overall microbial parameters | Prokaryote clusters | Fungal clusters | Putative metabolic traits and functions |
|---|---|---|---|---|---|---|
| Past | 2012 invasion effect size diversity | direct | - | - | ↑4 ↓8 | ↑ Saprotrophs and plant pathogens.<br>↓ Bacterial and fungal biomass; generalist soil saprotrophs. |
| | | via ↓NO2 | ↓bacterial biomass<br>↓fungal biomass | - | - | |
| Past | DCA1 trajectory | direct | - | ↓8 | ↓10 ↓11<br>↓13 | ↓ Specialist chitin-degraders; soil, wood, litter saprotrophs, plant pathogens and root endophytes, generalist algal parasites. |
| Past | DCA2 trajectory | direct | - | - | - | - |
| Past | **DCA3 trajectory** | direct | - | ↑4 | ↑4 ↓11<br>↓16 | ↑ Generalist chemoheterotrophs, various plant-phytopathogens; diverse group of generalist soil saprotrophs and plant pathogens.<br>↓ Specialist, ubiquitous soil and plant rhizosphere chemoheterotrophs; generalist algal parasites and unkown fungi. |
| | | **via ↑orgC** | - | ↓5 | **↑2** | |
| Past | **2012 invasion effect size DCA1** | direct | - | ↑8 | ↓12 | ↑ Specialist chitin-degraders; diverse group of generalist soil saprotrophs and plant pathogens.<br>↓ Specialist, ubiquitous soil and plant rhizosphere chemoheterotrophs; specialist litter saprotrophs and algal parasites. |
| | | **via ↑orgC** | - | ↓5 | **↑2** | |
| Past | **2012 invasion effect size DCA2** | **direct** | ↑prokaryote diversity | **↓1** ↑7 | **↑1** ↓8<br>↑14 ↓15 | ↑ Specialist, ubiquitous soil and plant rhizosphere bacteria, including methanotrophs and chitin-degraders; diverse group of generalist and specialist soil saprotrophs.<br>↓ Diverse group of generalist, ubiquitous soil and plant rhizosphere chemoheterotrophs; diverse group of generalist soil saprotrophs and plant pathogens; litter and dung saprotrophs. |
| | | **via ↓orgC** | - | ↑5 | **↓2** | |
| Past | **2012 invasion effect size DCA3** | **direct** | ↓prokaryote diversity<br>↓fungal diversity | **↑1** **↓3** ↓8<br>↑9 | ↓5 ↑8 | ↑ Diverse group of generalist ubiquitous soil and plant rhizosphere chemoheterotrophs, unknown specialist bacteria; generalist soil saprotrophs.<br>↓ Diverse group of ammonia oxidising archaea, organic matter degraders and N-fixing taxa; specialist chitin-degraders; diverse group of soil saprotrophs and mycoparasites. |

Pathways in bold indicate pathways with a relative contribution > 5% (see Fig 6). Clusters in bold indicate dominant clusters (see Fig 3). Clusters with a thick underline were classified as relative habitat generalists and clusters with a wave underline were classified as relative habitat specialists (see Fig S9). Clusters highlighted in green were enriched in natural grassland soil and clusters highlighted in red were enriched in abandoned arable soil (see Fig S10). For cluster specific information, see Table S3 and S5.

**Table S9** Sown and invaded plant species and their abbreviations

| Plant species | Abbreviation | Sown/invaded |
|---|---|---|
| *Acer spp* | Acesp | Invaded |
| *Agrimonia eupatorium* | Agreu | Sown |
| *Agrostis spp* | Agrsp | Invaded |
| *Anthericum ramosum* | Antra | Sown, not established |
| *Anthylis vulneraria* | Antvu | Sown |
| *Arabidopsis thaliana* | Arath | Invaded |
| *Arenaria serpyllifolia* | Arese | Invaded |
| *Arrhenatherum elatior* | Arrel | Invaded |
| *Artemisia vulgaris* | Artvu | Invaded |
| *Asperula spp* | Aspsp | Sown |
| *Aster amellus* | Astam | Sown, not established |
| *Astragalus cicer* | Astci | Sown |
| *Astragalus glycyphylos* | Astgl | Sown |
| *Atriplex spp* | Atrsp | Invaded |
| *Brachypodium pinnatum* | Brapi | Sown |
| *Bromus erectus* | Broer | Sown |
| *Bromus mollis* | Bromo | Invaded |
| *Bupleurum falcatum* | Bupfa | Sown |
| *Calamagrostis epigejos* | Calep | Invaded |
| *Campanula gentilis* | Camge | Sown |
| *Campanula glomerata* | Camgl | Sown |
| *Campanula patula* | Campa | Invaded |
| *Carex flacca* | Carfl | Sown |
| *Carex hirta* | Carhi | Invaded |
| *Cardamine spp* | Carsp | Invaded |
| *Carex tomentosa* | Carto | Sown |
| *Centaurea jacea* | Cenja | Sown |
| *Centaurea scabiosa* | Censc | Sown |
| *Cerastium holosteoides* | Cerho | Invaded |
| *Cirsium acaule* | Cirac | Sown, not established |
| *Cirsium pannonicum* | Cirpa | Sown |
| *Conyza spp* | Consp | Invaded |
| *Coronila varia* | Corva | Sown, not established |
| *Crepis biennis* | Crebi | Invaded |
| *Dactylis glomerata* | Dacgl | Invaded |
| *Daucus carota* | Dauca | Invaded |
| *Dianthus carthusianorum* | Diaca | Sown, not established |
| *Dipsacus sylvestris* | Dipsy | Invaded |
| *Elymus repens* | Elyre | Invaded |
| *Epilobium spp* | Episp | Invaded |
| *Erigeron annuus* | Erian | Invaded |
| *Euphorbia cyparissias* | Eupcy | Invaded |
| *Fallopia convolvulus* | Falco | Invaded |
| *Falcaria vulgaris* | Falvu | Invaded |
| *Galeopsis spp* | Galsp | Invaded |
| *Geranium sibiricum* | Gersi | Invaded |
| *Helianthemum grandiflorum* | Helgr | Sown |
| *Heracleum mantegazzianum* | Herma | Invaded |
| *Hieracium spp* | Hiesp | Invaded |
| *Holcus mollis* | Holmo | Invaded |
| *Hypericum perforatum* | Hyppe | Invaded |
| *Inula hirta* | Inuhi | Sown |
| *Inula salicina* | Inusa | Sown |

**Table S9** Continued

| Plant species | Abbreviation | Sown/invaded |
|---|---|---|
| *Lactuca serriola* | Lacse | Invaded |
| *Lamium purpureum* | Lampu | Invaded |
| *Laserpitium latifolium* | Lasla | Sown |
| *Lathyrus pratensis* | Latpr | Invaded |
| *Leontodon autumnalis* | Leoau | Invaded |
| *Leontodon hispidus* | Leohi | Sown |
| *Linum catharticum* | Linca | Invaded |
| *Linum flavum* | Linfl | Sown |
| *Linum tenuifolium* | Linte | Sown |
| *Lolium perenne* | Lolpe | Invaded |
| *Lotus corniculatus* | Lotco | Sown |
| *Medicago falcata* | Medfa | Sown |
| *Medicago lupulina* | Medlu | Invaded |
| *Myos spp* | Myosp | Invaded |
| *Plantago lanceolata* | Plala | Invaded |
| *Plantago media* | Plame | Sown |
| *Poa annua* | Poaan | Invaded |
| *Poa trivialis* | Poatr | Invaded |
| *Polygonum spp* | Polsp | Invaded |
| *Primula veris* | Prive | Sown, not established |
| *Prunella grandiflora* | Prugr | Sown |
| *Ranunculus spp* | Ransp | Invaded |
| *Raphanus raphanistrum* | Rapra | Invaded |
| *Rumex spp* | Rumsp | Invaded |
| *Salvia pratensis* | Salpr | Sown |
| *Salix spp* | Salsp | Invaded |
| *Salvia verticilata* | Salve | Sown |
| *Sanquisorba minor* | Sanmi | Sown |
| *Scabiosa ochroleuca* | Scaoc | Sown |
| *Silene vulgaris* | Silvu | Invaded |
| *Solidago canadensis* | Solca | Invaded |
| *Sonchus spp* | Sonsp | Invaded |
| *Stachys recta* | Stare | Sown |
| *Stipa spp* | Stisp | Invaded |
| *Tanacetum corymbosum* | Tanco | Sown |
| *Tanacetum spp* | Tansp | Invaded |
| *Tanacetum vulgaris* | Tanvu | Invaded |
| *Taraxacum officinalis* | Tarof | Invaded |
| *Teucrium chamaedrys* | Teuch | Sown |
| *Thlaspi arvense* | Thlar | Invaded |
| *Thymus pulegioides* | Thypu | Sown |
| *Trifolium medium* | Trime | Sown |
| *Trifolium montanum* | Trimo | Sown |
| *Trifolium pratensis* | Tripr | Invaded |
| *Trifolium repens* | Trire | Invaded |
| *Tussilago farfara* | Tusfa | Invaded |
| *Urtica dioica* | Urtdi | Invaded |
| *Veronica teucrium* | Verteu | Sown |
| *Viccia spp* | Vicsp | Invaded |
